# Systems genetics uncovers microbe-lipid-host connections in the murine gut

**DOI:** 10.1101/2021.11.29.470403

**Authors:** Q Zhang, V Linke, KA Overmyer, LL Traeger, K Kasahara, IJ Miller, DE Manson, TJ Polaske, RL Kerby, JH Kemis, EA Trujillo, TR Reddy, JD Russell, KL Schueler, DS Stapleton, ME Rabaglia, M Seldin, DM Gatti, GR Keele, DT Pham, JP Gerdt, EI Vivas, AJ Lusis, MP Keller, GA Churchill, HE Blackwell, KW Broman, AD Attie, JJ Coon, FE Rey

## Abstract

The molecular bases of how host genetic variation impact gut microbiome remain largely unknown. Here, we used a genetically diverse mouse population and systems genetics strategies to identify interactions between molecular phenotypes, including microbial functions, intestinal transcripts and cecal lipids that influence microbe-host dynamics. Quantitative trait loci (QTL) analysis identified genomic regions associated with variations in bacterial taxa, bacterial functions, including motility, sporulation and lipopolysaccharide production, and levels of bacterial- and host-derived lipids. We found overlapping QTL for the abundance of *Akkermansia muciniphila* and cecal levels of ornithine lipids (OL). Follow-up studies revealed that *A. muciniphila* is a major source of these lipids in the gut, provided evidence that OL have immunomodulatory effects and identified intestinal transcripts co-regulated with these traits. Collectively, these results suggest that OL are key players in *A. muciniphila-*host interactions and support the role of host genetics as a determinant of responses to gut microbes.

## Introduction

The gut microbiome plays fundamental roles in mammalian physiology and human health^1–3^. Environmental exposures, including maternal seeding, diet and cohabitation, and host genetic variation modulate gut microbiota composition^4–6^ and contribute to the large degree of interpersonal variation observed in human gut microbial communities. These inter-individual differences have been associated with disparate responses to diet, pathogens and therapeutic drugs^7–10^. Understanding how host genetics and environmental factors interact to shape the composition of the gut microbiota is imperative for designing health-promoting strategies aimed at modifying its composition.

Recent advances in high throughput sequencing have fueled progress in our understanding of the impact of host genetics and the gut microbiome on health. Genome-wide association studies (GWAS) have revealed host genetic-gut microbial trait associations in human cohorts^11–15^. Furthermore, microbiome-wide association studies leveraging host genetic information and Mendelian randomization have highlighted connections between the gut microbiome and other molecular complex traits including fecal levels of short-chain fatty acids^16^, plasma proteins^17^, and ABO histo-blood group type^18^ in humans. The effects of host genetic variation on gut microbiome composition have also been previously examined in different mouse populations^19, 20^. However, most of these studies have focused on microbial organismal composition and there is currently a major gap in our understanding of the impact of host genetic variation on the functional capacity of the gut microbiome.

Microbial metabolites are critical nodes of communication between microbes and the host. These include small molecules derived from dietary components (e.g., TMAO)^21^, or *de novo* synthesized by microbes such as vitamins^22^ and lipids^23^. Lipids including eicosanoids, phospholipids, sphingolipids and fatty acids act as signaling molecules to control many cellular processes^24–26^. Gut microbes not only modulate absorption of dietary lipids via regulation of bile acid production and metabolism but also are a major source of lipids and precursor metabolites for lipids produced by the host^27, 28^. Bacterial cell membrane-associated lipids are also important for microbe-host interactions^23, 29^, although our understanding of their roles in these dynamics is only emerging for gut bacteria.

Defining the general principles that govern microbe-host interactions in the gut ecosystem is a daunting task. Systems genetic studies can generate novel hypotheses that invoke processes and molecules that have no precedent, which can be used for the identification of genes, pathways and networks underlying these interactions. To investigate the connections between gut microbes, intestinal lipids and host genetic variation, we leveraged the Diversity Outbred (DO) mouse cohort, a genetically diverse population derived from eight founder strains: C57BL6J (B6), A/J (A/J), 129S1/SvImJ (129), NOD/ShiLtJ (NOD), NZO/HLtJ (NZO), CAST/EiJ (CAST), PWK/PhJ (PWK), and WSB/EiJ (WSB)^30, 31^. These eight strains harbor distinct gut microbial communities and exhibit disparate metabolic responses to diet-induced metabolic disease^7^. The DO population is maintained by an outbreeding strategy aimed at maximizing the power and resolution of genetic mapping. We characterized the fecal metagenome, intestinal transcriptome and cecal lipidome in DO mice and performed QTL analysis to identify host genetic loci associated with these traits. We integrated microbiome QTL (mbQTL) and cecum lipidome QTL (clQTL) to uncover novel microbe-lipid associations and identified candidate genes expressed in the distal small intestine associated with these co-mapping traits. These datasets represent a valuable resource for interrogating the molecular mechanisms underpinning interactions between the host and the gut microbiome.

## Results

### Gut microbial features are associated with host genetics

We characterized the fecal microbiome from 264 DO mice fed a High-Fat High-Sucrose (HF/HS) diet for ∼22 weeks (Fig. 1). Metagenomic analysis revealed ∼1.9 million unique predicted microbial open reading frames (i.e., metagenes). Annotation of non-redundant (NR) metagenes against the Kyoto Encyclopedia of Genes and Genomes (KEGG) orthology database and taxonomic assignment against the NCBI database identified 2803 functional orthologs (i.e., KOs) and 187 bacterial taxa across all mice. We also performed metagenomic binning to obtain metagenome-assembled genomes (MAGs), corresponding to species-level bacterial genomes. (**Supplementary Fig. 1**, **Supplementary Table 1**-4; see Supplementary Notes for details on metagenomic analysis).

**Figure 1.**
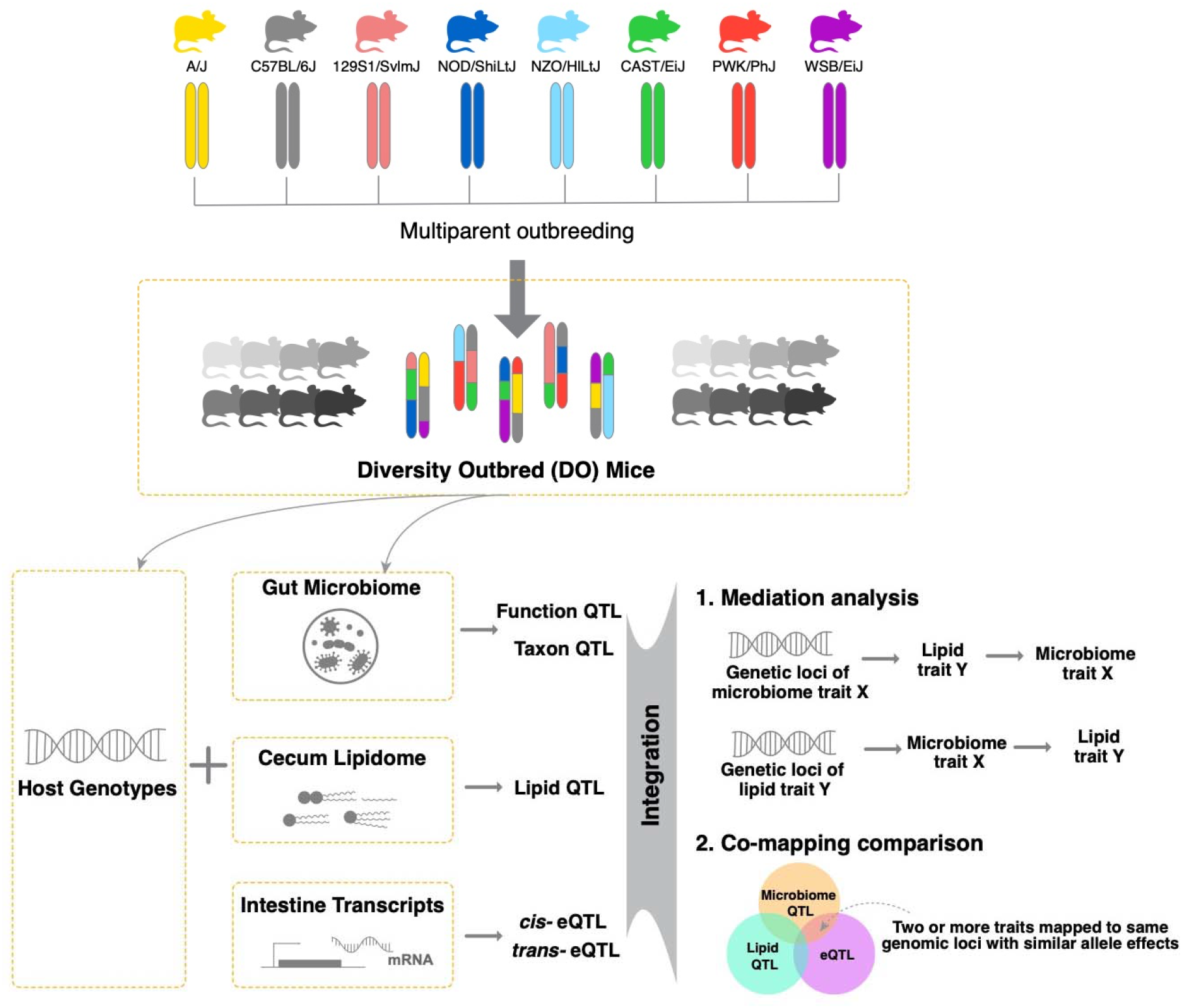
Overview of the study. Fecal metagenomes (n=264), cecal lipidomes (n=381) and distal small intestine transcriptomes (n=234) were generated from Diversity Outbred mice. Quantitative trait loci (QTL) analysis identified genomic regions associated with variations in bacterial taxa, bacterial functions, levels of bacterial- and host-derived lipids and small intestine transcript levels. Mediation analysis and co-mapping comparisons were used to identify causal links between traits.

We next used QTL analysis to identify regions of the mouse genome associated with the abundance of these traits. We detected 2814 suggestive associations for KOs (LOD > 6, *P*_Genome-wide-adj_ < 0.2), and 200 suggestive associations for bacterial taxa (LOD > 6*, P*_Genome-wide-adj_ < 0.2). 15 genomic loci were highly associated with microbial KOs (LOD >8.97, *P*_Genome-wide-adj_ < 0.05) and 7 genomic loci were highly associated with bacterial taxa (LOD >7.92, *P*_Genome-wide-adj_ < 0.05) (Fig. 2a, **Supplementary Table 5**-6). A particularly striking finding among these genomic loci was the presence of QTL “hotspots” where genomic regions were associated with multiple microbial traits (**Supplementary Table 7**). Among these, the hotspot on chromosome 15 at ∼63.5 Mbp was the largest, where we identified 154 microbial traits with LOD score > 6. We estimated DO founder allele effects as best linear unbiased predictors (BLUP) for the traits that mapped to this locus. Among these we detected two clear groups of traits that exhibited opposite allele effects: a group of KOs and taxa showing positive association with the 129 allele, and another group of KOs and taxa that were negatively associated with the 129 allele (**Supplementary Fig. 2a**). As detailed below, the two most abundant gut bacterial phyla, Firmicutes and Bacteroidetes, mapped to this locus with opposite allele effects.

**Figure 2.**
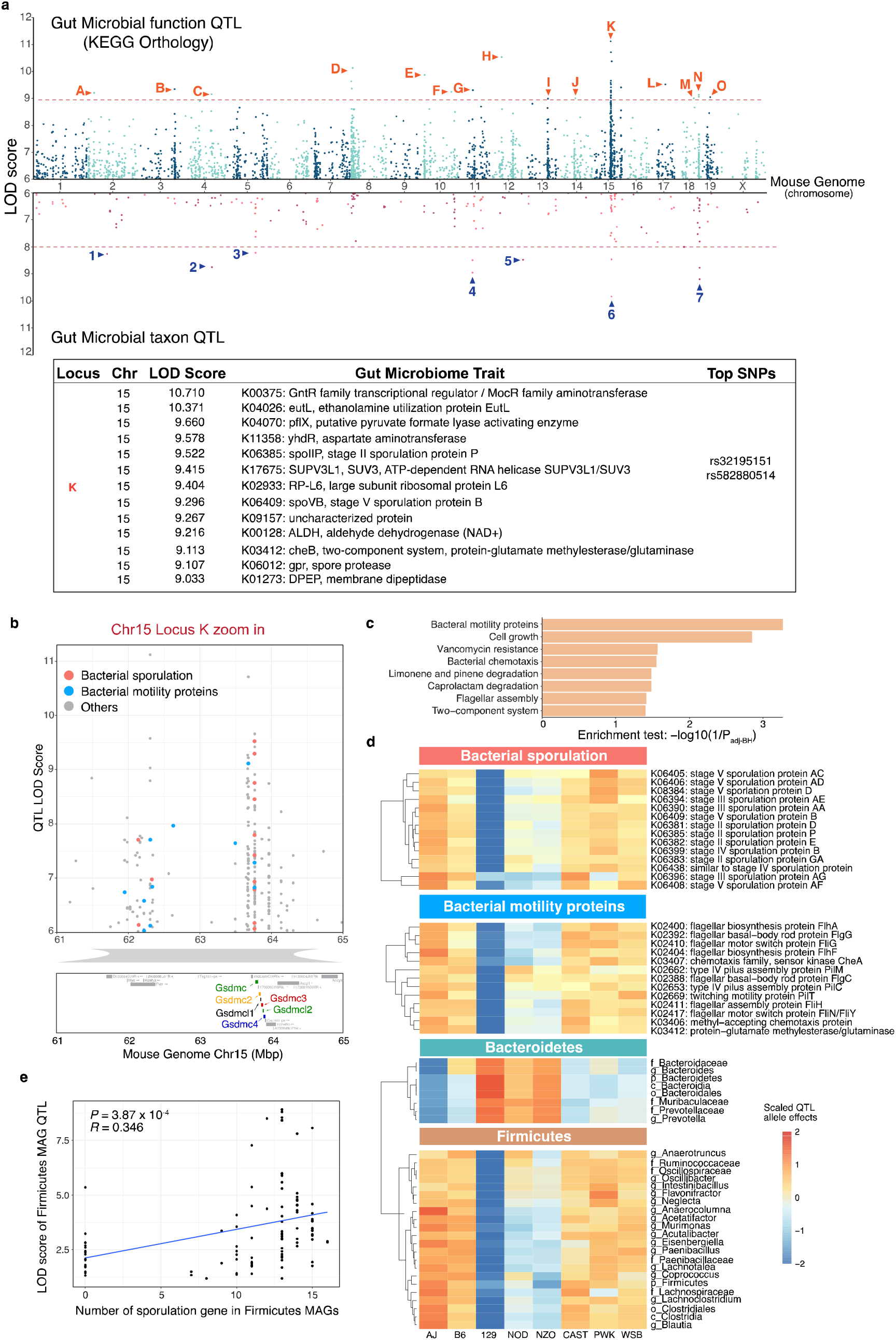
Genetic architecture of quantitative trait loci (QTL) for microbial traits in the Diversity Outbred (DO) mouse cohort. **a,** QTL mapping results for 2803 gut microbial KEGG Orthology function traits (KO) and 187 bacterial taxa traits, using sex, days in diet, and cohort as covariates. Each dot represents a QTL on the mouse genome for a given trait. Significance thresholds for QTL “hostspots” were determined by permutation tests (LOD > 8.97, *P*_Genome-wide-adj_ < 0.05 for microbial function traits; LOD > 7.92, *P*_Genome-wide-adj_ < 0.05 for bacterial taxon traits). Capital letters indicate highlighted loci where multiple microbial function traits map; numbers indicate highlighted loci where significant associations with taxa are observed. The gut microbiome traits that mapped to these loci are listed in **Supplementary Table 3**. **b,** Gut microbiome QTL hotspot on Chr15 has multiple bacterial sporulation and motility functions mapping to it. **c,** Enrichment analysis for functions mapping at hotspot on Chr15**. d**, QTL for microbial functions that mapped to Chr15 hotspot had negative 129 allele effects. QTL for Firmicutes mapping to Chr15 hotspot had negative 129 allele effects whereas QTL for Bacteroidetes mapping to this locus had positive 129 allele effects. **e,** Correlation analysis between the number of sporulation KOs detected in Firmicutes MAGs mapping at Chr15 QTL hotspot and the LOD scores for these MAGs (*P* = 3.87 x 10^-3^, *R* = 0.346).

Pathway enrichment analysis showed that bacterial “motility proteins” and “cell growth” functional categories were significantly enriched in the group of KOs associated most strongly with 129 alleles (Fig. 2b-c). More specifically, abundances of 14 sporulation functions were negatively associated with 129 alleles (Fig. 2d). Further investigation of the KO distribution across all MAGs revealed that all bacterial sporulation KOs were only present in MAGs belonging to Firmicutes, whereas most of KOs that showed positive 129 allele effects were present in MAGs belonging to Bacteroidetes (**Supplementary Fig. 2b**). To assess whether the allele effects observed from QTL mapping corresponded to the trait patterns in the DO founder strains, we examined previously published 16S rRNA gene data from age-matched mice from the eight founder strains, also fed a HF/HS diet^20^. Consistent with these findings we found that the 129 mouse strain had higher levels of Bacteroidetes and the highest Bacteroidetes/Firmicutes ratio (**Supplementary Fig. 2c**). Interestingly, we detected a significant positive correlation between the number of sporulation KOs in Firmicutes MAGs mapping at this locus and the LOD scores for these MAGs (Fig. 2e). Importantly, Firmicutes MAGs commonly detected in our dataset that do not contain sporulation KOs (e.g., *Lactobacillus*, *Lactococcus*) did not exhibit significant association to this QTL. These results support the notion that host genetic variation affects gut community structure in part by modulating the abundance of sporulating bacteria.

Single nucleotide polymorphism (SNP) association analysis within the Chr15 QTL hotspot identified six significant SNPs: two intron variants, SNP rs582880514 in the *Gsdmc* gene and SNP rs31810445 in the *Gsdmc2* gene, both with LOD scores of 8.0. The other four SNPs were intergenic variants (**Supplementary Fig. 2d**). Gasdermins (Gsdm) are a family of pore-forming proteins that cause membrane permeabilization and pyroptosis^32^, an inflammatory form of programmed cell death that is triggered by intra- and extracellular pathogens^33^. These results indicate that host genetic variation in *Gsdmc*/*Gsdmc2* genes is associated with abundance of gut bacterial functions and raises the hypothesis that these host proteins could modulate the abundance of bacterial taxa harboring motility and/or sporulation functions.

### Cecal lipid features are associated with gut microbes and host genetics

We employed a broad discovery strategy to agnostically detect lipid actors potentially relevant to gut microbiome-host interactions. We used liquid chromatography coupled with tandem mass spectrometry (LC-MS/MS) to characterize the cecal lipidome of 381 DO mice, including all mice used for the metagenomic analysis. We identified 1,048 lipid species representing 35 lipid classes (Fig. 3a-b) and the four major lipid categories: (i) fatty acyls, (ii) phospholipids, (iii) sphingolipids, and (iv) glycerolipids. The highest numbers of lipids were recorded for the classes of triglycerides (TG) and phosphatidylcholines (PC), species known to be abundant in the mammalian host^34^. Due to the nature of the cecal sample, we can assume that we observed lipids derived from diet, host and microbes. Phosphatidylglycerols (PG), for example, which represent the phospholipid class with the second most members in our data, are known to be a major component in the bacterial lipidome^35^. In mammals, on the other hand, PG are only a minor component. Similarly, among glycerolipids, monogalactosyldiacylglycerols (MGDG) account for the second highest number lipids detected in this class. While found at high levels in bacteria and plants, these lipids are only minor components of animal tissue^36^. These findings suggest that the cecal lipidome include components of the host and the gut microbiome. Correlation analysis between MAGs and cecal lipids abundance, plus comparison of the cecal lipidome of conventionally-raised *vs.* germ-free mice identified taxa that potentially modulate abundance of lipids in the gut. (Supplementary notes and **Supplementary Fig. 3**, **Supplementary Table 8**-10). Furthermore, QTL mapping identified 457 significant QTL for cecal lipid features (LOD > 7.5, *P*_Genome-wide-adj_ < 0.05) (Supplementary notes and Fig. 3c, **Supplementary Table 11**). Altogether these associations provide a wealth of information offering potential molecular descriptors of the genetic regulation of the microbiome.

**Figure 3.**
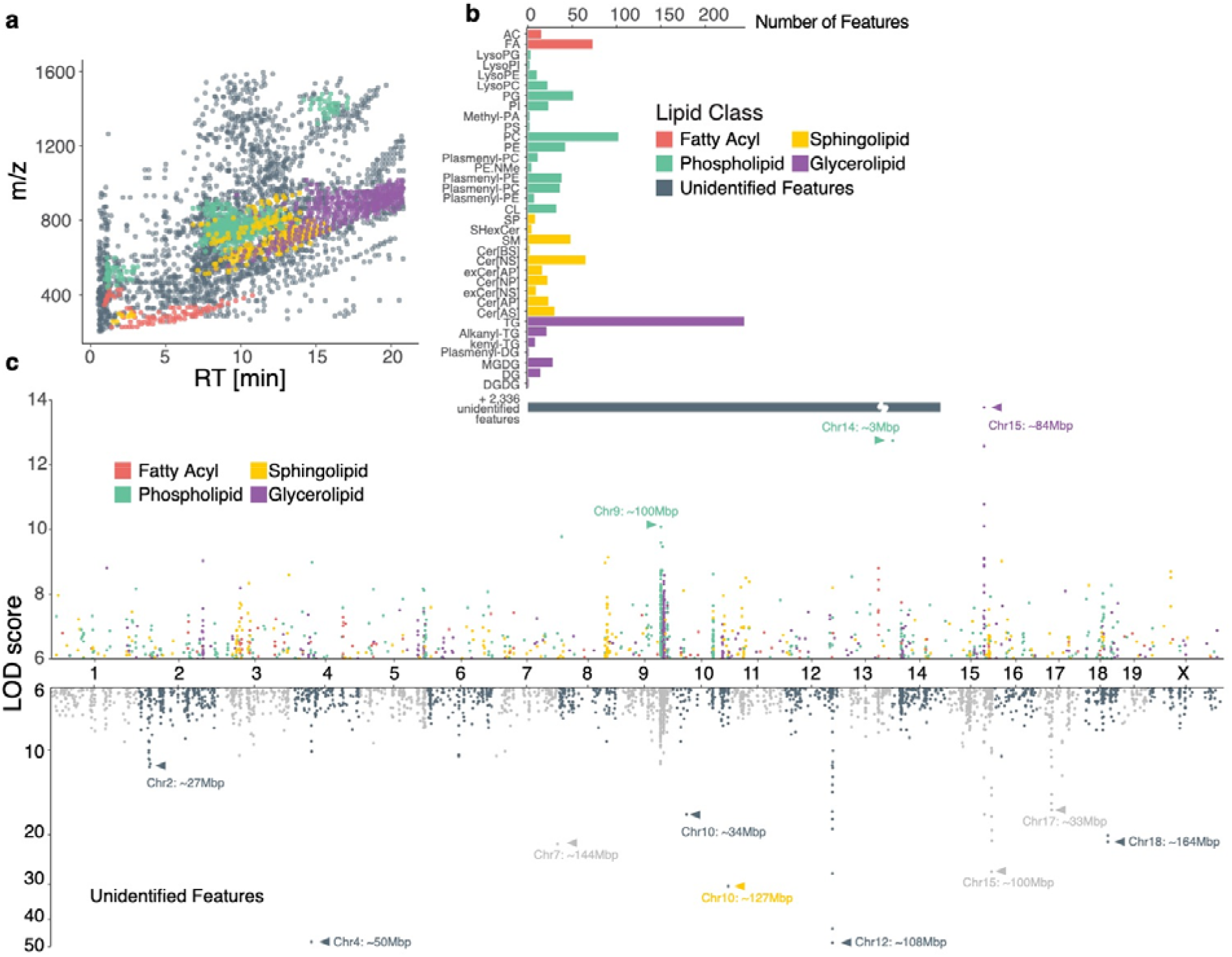
Genetic architecture of the cecal lipidome in Diversity Outbred (DO) mice. **a**, A total of 3,384 cecum lipid features were quantified across 381 DO mice, 1,048 of which were identified as lipids from four major classes. Features of each class occupied characteristic regions in the m/z – RT space. **b**, Identified lipids belonged to 35 lipid subclasses, with bacteria-associated phosphatidylglycerol (PG) and monogalactosyldiacylglycerol (MGDG) as common subclasses. **c,** 3,964 suggestive cecal lipid QTL (LOD > 6, *P*_Genome-wide-adj_ < 0.2) and 12 QTL hotspots were identified; 68.2% of the identified lipids showed a total of 1,162 QTL while a similar portion of 70.1% of unidentified features contributed 2,802 QTL. For lipid class abbreviations see **Supplementary Table 16**. RT = retention time.

### Mediation analysis reveals link between *Akkermansia muciniphila* and cecal lipid features

To identify causal links between gut microbial traits and cecal lipid traits, we performed mediation analysis between individual gut microbial metagenes and lipid features that co-map (Methods). Mediation analysis seeks to determine whether a QTL has separate effects on two traits, or if it affects one trait through its effect on another trait, in which case the intermediate trait is called a mediator. Figure 4a shows gut microbial metagenes mediating the QTL effect on a cecal lipid trait. We reasoned that if a microbial trait had an effect on a cecal lipid that was independent from the cecal lipid’s QTL, its inclusion as a covariate would be unlikely to affect the cecal lipid QTL signal significantly. However, for microbial traits that mediate the QTL effect on the cecal lipid, there would be a large drop in the original cecal lipid QTL LOD score. Interestingly, we found three cecal lipid features with QTL that were mediated by microbial metagenes. Most of these mediating microbial traits were genes belonging to the bacterium *A. muciniphila*. It is important to note that the direction of the causal effect between microbial trait and cecal lipid cannot be directly inferred from the data, but these results suggest *A. muciniphila* levels and the abundance of these lipid species in the gut are modulated by the same host genetic variation and that the traits are potentially causally related.

**Figure 4.**
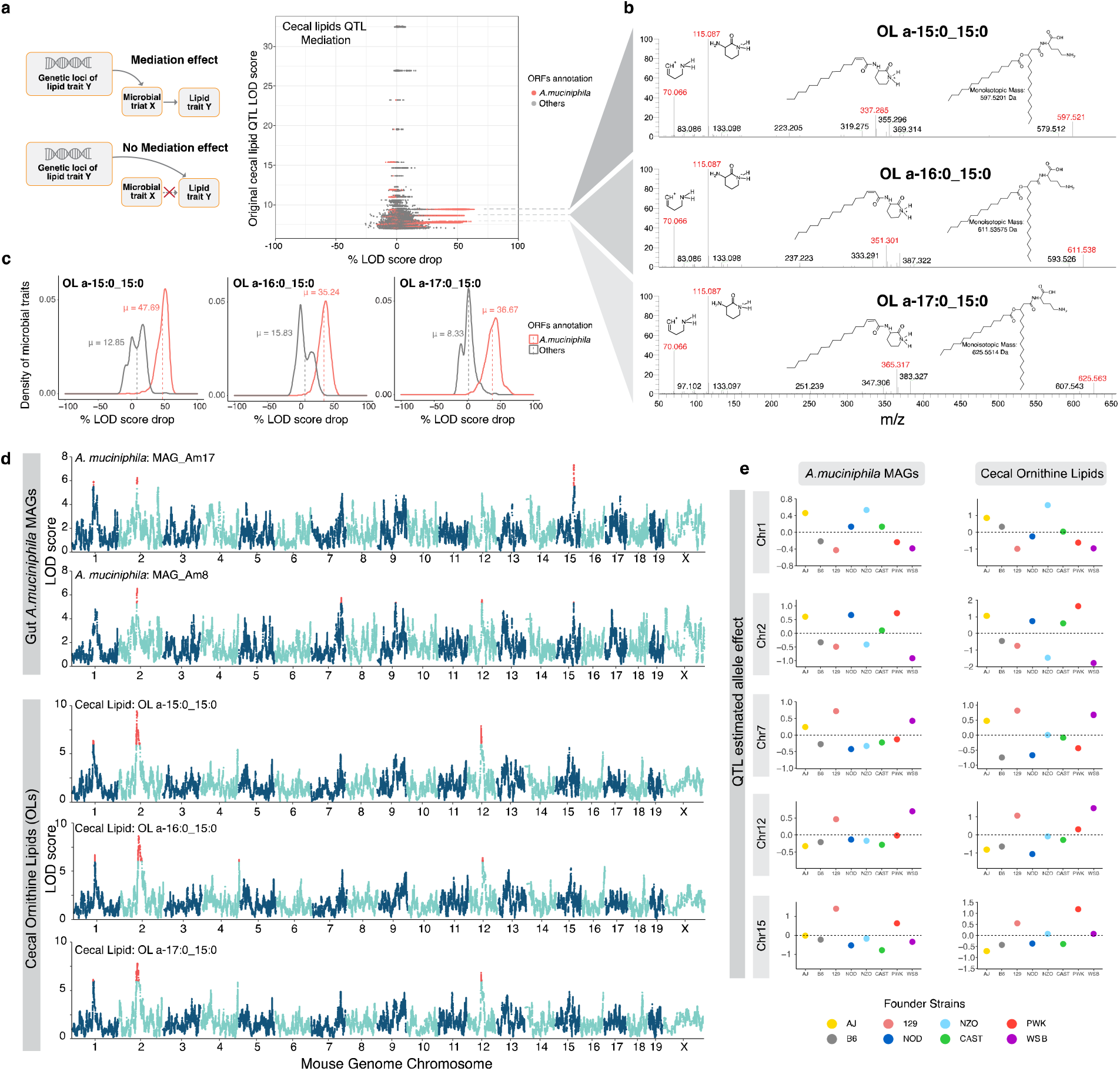
Mediation analysis revealed potential causal relationship between *A. muciniphila* and ornithine lipids (OL). **a,** Drop in QTL LOD score for cecal lipid features when individual gut microbial metagenes are added as covariates. Association of three unknown cecal lipid features with host genome is impacted by *A. muciniphila* genes. **b,** The three lipid features mediated by *A. muciniphila* genes were identified as OL (insert shows OL structure). **c,** Distribution of three OL LOD score drop when adding individual gut *A. muciniphila* genes as covariates (Mediation model) or adding individual genes not from gut *A. muciniphila* as covariates (Null model). **d,** Three OL species QTL co-mapped at five loci (Chr1, Chr2, Chr7, Chr12, Chr15) with *A. muciniphila* MAGs QTL. **e,** Founder allele effects for *A. muciniphila* MAGs and cecal OL were estimated in the DO population from the founder strain coefficients observed at the corresponding QTL at each locus.

We further tested whether the cecal lipids and *A. muciniphila* mapped to the same loci as expected. Mapping of the 46 reconstructed *A. muciniphila* MAGs to the host genome revealed multiple QTL including Chr1: 92.9Mbp, Chr2: 79.4Mbp, Chr7: 129.8Mbp, Chr12: 59.4Mbp, and Chr15: 75.9Mbp (Fig. 4d). Interestingly, the three cecal lipids also showed QTL at the same loci and exhibited similar founder allele effect patterns (Fig. 4e). These founder allele effects on *A. muciniphila* abundance are consistent with a previous study of gut bacterial abundance in the DO founder strains^20^. Although these lipid features were not initially identified by our lipidomic analysis pipeline, they appeared to be closely related to each other. Further analysis of their fragmentation spectra suggested that these unidentified features were ornithine lipids (OL) (Fig. 4b-c and Supplementary Notes). This was confirmed with a synthetic OL (see below). The three features would have the sum compositions of OL 30:0, OL 31:0, and OL 32:0, detected as [M+H]+ ions. In OL, a 3-hydroxy fatty acid is connected via an amide linkage to the ornithine amino acid that serves as the headgroup. A second fatty acid is then connected to the first via an ester linkage^37^. OL are bacteria-specific non-phosphorus glycolipids that are found in the outer membranes of selected Gram-negative bacteria^38, 39^.

### *A. muciniphila* produces OL in the mouse and human gut

*A. muciniphila* is a Gram-negative bacterium that has been associated with many metabolic host benefits^40, 41^. Several Gram-negative bacteria are known to produce OL under phosphate-limited conditions^39^. While previous research suggests that OL are important for microbe-host interactions in rhizobiales^42^ and

*Pseudomonas aeruginosa*^29^, the occurrence of these lipids in gut bacteria was not known. To test whether *A. muciniphila* produces OL, we performed two validation experiments. First, we leveraged the discovery lipidomics LC-MS/MS assay described above to profile lipids in *A. muciniphila* and two other gram-negative species, *Bacteroides thetaiotaomicron* and *Escherichia coli* grown under anaerobic conditions. We found similarly high levels of all three targeted OL species in extracts from *A. muciniphila* but not in the other species, which were indistinguishable from the solvent blank (Fig. 5a). Since phosphate limitation triggers production of OL in some bacterial species^29^, in follow-up experiments we tested whether phosphate levels modulated abundance of OL in *A. muciniphila* grown *in vitro.* We examined three different levels of phosphate [0.02 mM (growth limiting), 0.2 mM (adequate) and 2 mM (excess)]. LC-MS/MS analysis confirmed that OL are a dominant lipid species detected in *A. muciniphila* cell extracts regardless of the phosphate levels included in the growth media (**Supplementary Fig. 4a-b**). Furthermore, OL were detected in extracellular vesicles isolated from *A. muciniphila*-grown *in vitro*, (**Supplementary notes** and **Supplementary Fig. 4c**). These results suggested that OL are likely localized in the *A. muciniphila* outer membranes and provide insights into how these lipids may interact with the host.

**Figure 5.**
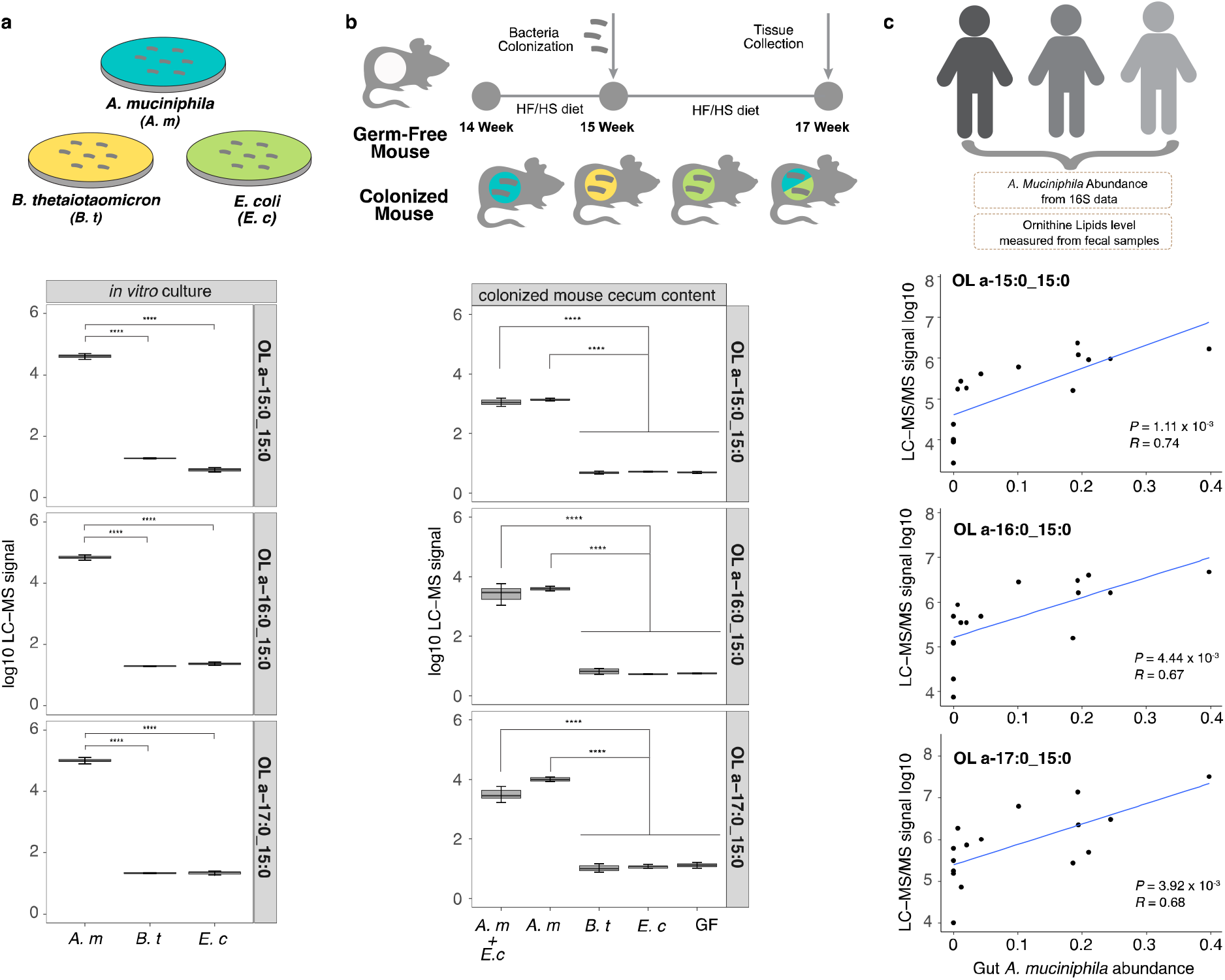
*A. muciniphila* produces OL in the mouse and human gut. **a,** OL abundance for the three major species detected in mice in cell pellets collected from *A. muciniphila* (*A. m*), *B. thetalotamicron* (*B. t*), and *E. coli* (*E. c*) grown *in vitro*. **b,** OL detected in cecal contents from gnotobiotic mice colonized with (i) *A. muciniphila,* (ii) *B. thetaiotaomicron,* (iii) *E. coli* and (iv) *A. muciniphila* plus *E. coli* for 2 weeks (n=3-4 mice/treatment). **c,** Detection of prominent OL species in human fecal samples is highly correlated with *A. muciniphila* abundance by correlation. Data are presented as mean ± SEM; ** *P* < 0.01, *** *P* < 0.001, **** *P* < 0.0001.

We further profiled lipids produced by *A. muciniphila* colonizing the gut of gnotobiotic mice. Five groups of adult germ-free B6 mice were mono-colonized with each of the species mentioned above, bi-associated with *E. coli* and *A. muciniphila* or kept germ free (n=3-5/group). Mice were maintained in the same HF/HS diet used for the DO study for two weeks after inoculation. LC-MS/MS analysis of cecal contents from these mice showed that only mice colonized with *A. muciniphila* had detectable levels of OL in their cecum (Fig. 5b). Altogether these results confirm that *A. muciniphila* gut colonization is causally linked with high levels of OL.

We examined whether *A. muciniphila* colonization is associated with the presence of OL in the human gut. We analyzed lipid content in fecal samples from a previously characterized cohort of old adults^43^; we selected samples with (i) low or no detectable levels of *A. muciniphila*, (ii) intermediate abundance of *A. muciniphila* (0.7% - 10%) and (iii) the highest levels of *A. muciniphila* detected (18.6% - 39.8%) in this human cohort. LC-MS/MS analysis of human fecal samples detected a broader range of OL species than axenic cultures or mice colonized with *A. muciniphila*, but the levels of the three previously identified OL 15:0_15:0, OL 16:0_15:0, and OL 17:0_15:0 were all significantly correlated with *A. muciniphila* levels (Fig. 5c). Together these results suggest that *A. muciniphila* is a major producer of OL in the mouse and human gut.

### OL modulate LPS-induced cytokines from bone marrow-derived macrophages (BMDM)

To test whether *A. muciniphila*-derived OL elicit immune responses on the host, we first chemically synthesized the most abundant OL detected in the DO mouse gut, i.e., OL_15:0_15:0 (**Supplementary Notes**). Because of the generally beneficial effects of *A. muciniphila* on host health previously documented in both human and mouse studies, and based on the structural similarity between OL and lipid A from LPS, we speculated that the OL might function as antagonists of lipid A. We examined the effects of the OL preparation in the absence and presence of LPS on cytokine production by bone-marrow-derived-macrophages (BMDM). Treatment with LPS induced a significant increase in the production of TNF-α and IL6 by BMDM obtained from B6 and 129 mice (**Supplementary Fig. 5a**). In contrast, treatment with OL did not stimulate significant production of TNF-α and IL6 by these cells (**Supplementary Fig. 5b**) except for a modest increase at 500ng/ml and 1000ng/ml. However, we observed that pretreatment of macrophages with OL had an inhibitory effect on LPS-induced TNFα and IL6 in both B6 and 129 mice, without causing significant changes in cell viability (**Supplementary Fig. 5c**). These results suggested that *A. muciniphila*-derived OL can prevent LPS-induced inflammation response. Furthermore, we measured other cytokines secreted by LPS-treated BMDM and observed that the OL preparation inhibited the production of IL-1β, MCP-1, MIP-1α, GM-CSF, IL12 and RANTES (Fig. 6), although there were differences in the responses to LPS and OL as a function of BMDM genetic background. In addition, OL increased levels of anti-inflammatory cytokine IL-10 in these cells (Fig. 6), suggesting OL may modulate inflammation by altering the levels of both pro-inflammatory and anti-inflammatory cytokines. Interestingly, production of IL-12 in the presence of LPS was more than ten times higher in 129 mice than B6 mice, and OL had a larger inhibitory effect in these mice (Fig. 6). These results indicate that *A. muciniphila-*derived OL may influence host innate immune responses and their effects may vary as a function of host genetics.

**Figure 6.**
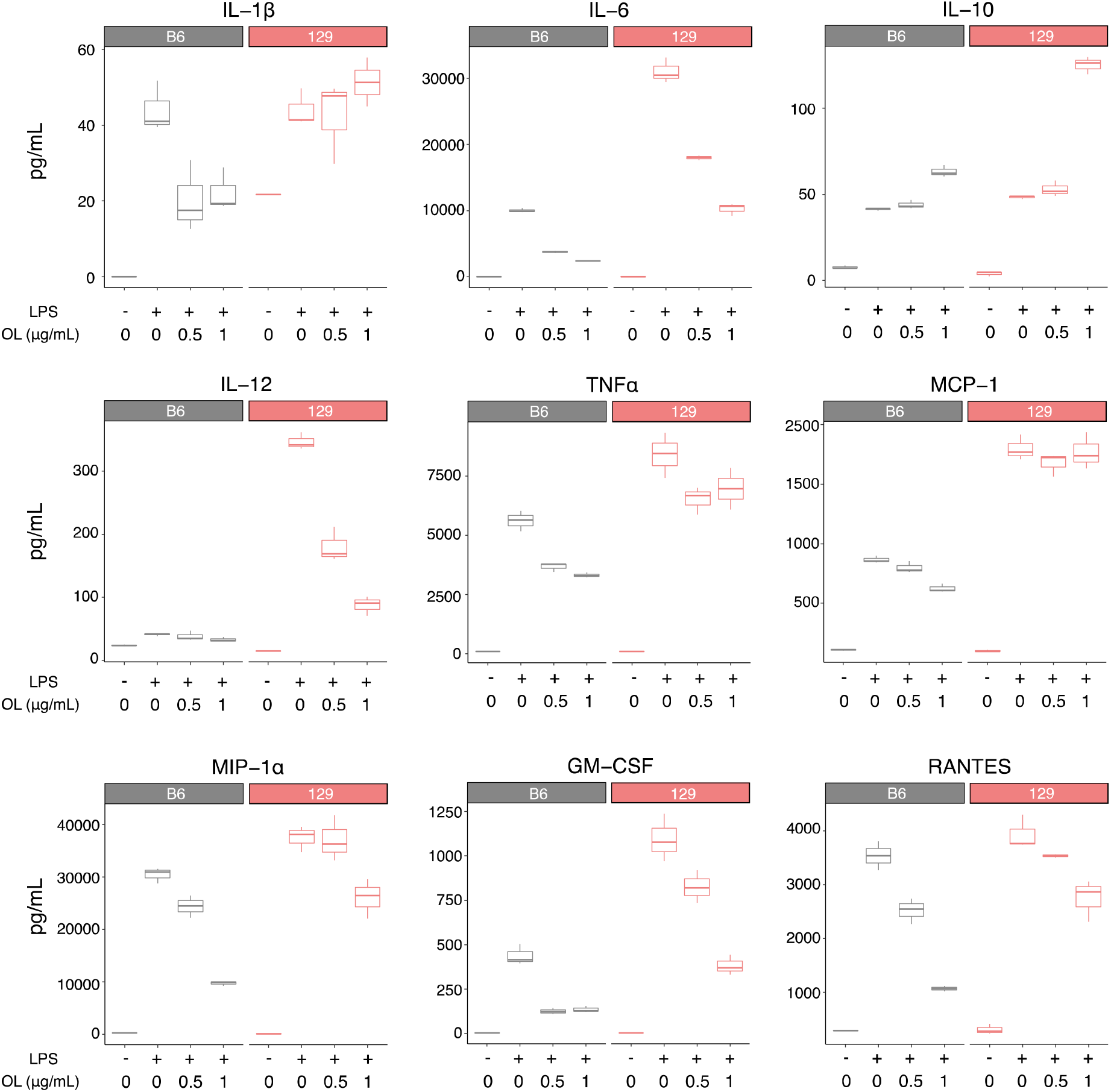
OL modulate LPS-induced production of cytokines from bone marrow-derived macrophages (BMDM). Levels of IL-1β, IL-6, IL-10, IL-12, TNF-α, MCP-1, MIP-1α, GM-CSF, and RANTES detected in supernatants from B6 and 129 mice BMDM stimulated with LPS (10 ng/ml) and different concentrations of OL.

### Expression QTL (eQTL) for intestinal innate immunity genes co-map with *A. muciniphila* and OL QTL

We profiled transcript levels in the distal small intestines of 234 DO mice using RNAseq. We detected 8,137 transcripts with a minimum of 10 counts/million (CPM) in at least 10% of DO mice. We identified 4,462 local eQTL with an average LOD score of 21.2 and 10,894 distal eQTL with an average LOD score of 7.1 (**Supplementary Table 12**). By comparing eQTL allele effects with those for the co-mapping mbQTL and clQTL, we identified gut microbial features and cecal lipids that were potentially co-regulated with intestinal transcripts (**Supplementary Fig. 6** and **Supplementary Notes**).

We searched the support intervals for the five co-mapping QTL regions for *A. muciniphila* and OL (Chr1, Chr2, Chr7, Chr12, and Chr15) for candidate host genes of interest using the eQTL data. By comparing the allele effects between co-mapping eQTL and the *A. muciniphila*/OL QTL, we identified several candidate host genes whose eQTL allele effects were correlated with *A. muciniphila*/OL (Fig. 7, **Supplementary Fig. 7, Supplementary Table 13**). At the Chr1 QTL region, there were four candidate genes: (i) Gene Activating transcription factor 3 (*Atf3*) had a distal eQTL at Chr1: 92.96Mbp with QTL LOD score of 6.55. At this locus AJ/NZO were the driver alleles, as is the case for the *A. muciniphila* and OL QTL. ATF3 plays an important role during host immune response events by negatively regulating the transcription of proinflammatory cytokines induced by the activation of toll-like receptor 4^44^. (ii) The gene TRAF-interacting protein with a forkhead-associated domain (*Tifa*), had a distal eQTL at Chr1: 90.95 Mbp with LOD score of 6.19 and also had the AJ/NZO driver alleles pattern. TIFA has been reported to sense bacterial-derived heptose-1,7-bisphosphate – an intermediate in the synthesis of LPS, via a cytosolic surveillance pathway triggering the NF-kB response^45, 46^. Additionally, TIFA interacts with TRAF6 to mediate host innate immune responses. (iii) The gene Jumonji domain-containing protein 8 (*Jmjd8*) had a distal eQTL at Chr1: 92.14Mbp with LOD score of 6.72. NZO is the driver allele, but it showed inverted allele effects compared to *A. muciniphila* and OL QTL. JMJD8 functions as a positive regulator of TNF-induced NF-kappaB signaling^47^. A recent study showed that JMJD8 was required for LPS-mediated inflammation and insulin resistance in adipocytes^48^. (iv) Gene *Gcg* had a distal eQTL at Chr1: 92.36Mbp with LOD score of 7.11. *Gcg* encodes multiple peptides including glucagon, glucagon-like peptide-1(GLP-1). GLP-1 levels are induced by a variety of inflammatory stimuli, including endotoxin, IL-1β and IL-6^49^. The finding that these genes with distal eQTL that co-map with *A. muciniphila* and OL QTL on Chr1 are involved in host immune responses to microbial-associated molecular patterns (MAMPs) such as LPS suggests that expression of these genes contribute to the regulation of host responses to OL and/or potentially modulate abundance of *A. muciniphila*.

**Figure 7.**
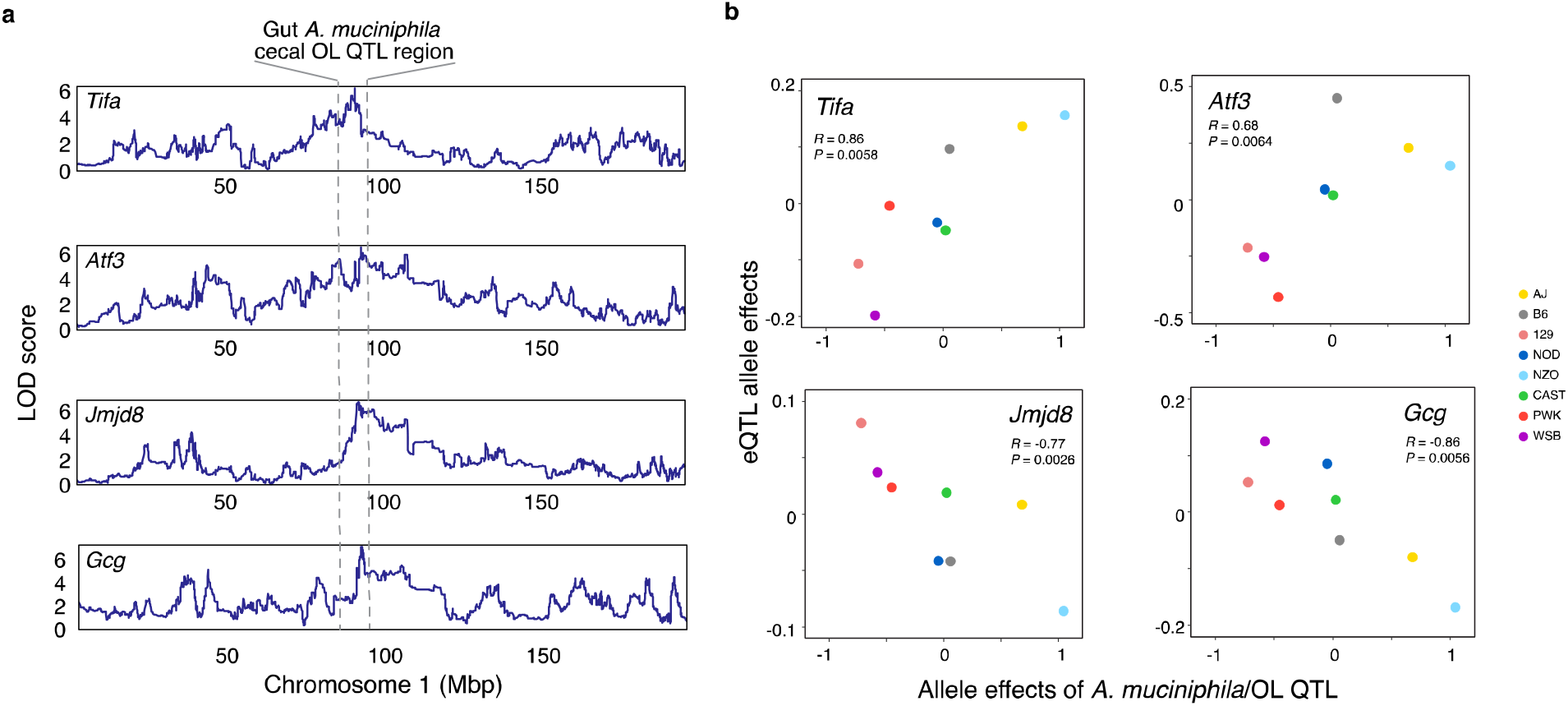
eQTL for distal small intestine (ileum) genes that co-map with *A. muciniphila* and cecal OL at chromosome 1. **a.** *A. muciniphila*, cecum OLs, and *Tifa*, *Atf3*, *Jmjd8* and *Gcg* co-map at Chr1: 90-95 Mbp. LOD score in Y axis represents significance of QTL for trait. **b.** Correlation of allele effects between candidate gene eQTL and *A. muciniphila*/OL QTL.

## Discussion

We applied a systems genetics approach to identify novel relationships between gut microbes, their encoded functions, cecal lipids and host intestinal gene expression. We identified bacterial functions influenced by host genetic variation and discovered that the bacterium *A. muciniphila* produces immunoactive OL that are detected in fecal samples from humans and mice colonized with this bacterium. *A. muciniphila* has been previously associated with host genetic variation in both mice and humans at several loci^15, 19, 50, 51^, however distinct with what we report here.

Previous work suggested that some Gram-negative bacteria produce OL under phosphate-limiting conditions^53–55^. In contrast we observed that OL levels were consistently high across a 100-fold phosphate level range, suggesting that phosphate is not a major driver of OL synthesis in *A. muciniphila*. Notably, a recent study showed that increased OL production by the bacterial pathogen *P. aeruginosa* makes its cellular surface more hydrophobic, and resulted in lower virulence, and higher resistance to antimicrobials and host immune defenses^29^. *A. muciniphila* consumes host glycans present in the mucus layer, which is in proximity to the host epithelium. While mucin carbohydrates and amino acids serve as substrates for *A. muciniphila,* there are also soluble host defense molecules trapped in this layer that prevent invasion of microbes to the underlying mucosal epithelial cells. We speculate that membrane OL impact interactions of *A. muciniphila* with the intestinal milieu and may represent an adaptation critical to its niche and important for its interactions with the host. Development of tools to genetically manipulate *A. muciniphila* will be needed to test these hypotheses.

The inhibitory effects of OL on LPS-induced cytokines that we and others have observed^56, 57^ may represent an important aspect of how *A. muciniphila* impact host physiology. Previous studies identified both natural and synthetic molecules that can inhibit TLR4-mediated LPS signaling; compounds that prevent septic shock, and have anti-inflammatory and anti-neuropathic pain activities *in vivo*^58^. One group of LPS antagonist molecules targeting CD14 share structural features to *A. muciniphila* OL including a glucose unit linked to two hydrophobic chains and a basic nitrogen on C-6^59^, supporting the potential anti-inflammatory effects of OL. Although the precise mechanisms of how OLs inhibit LPS signaling are unknown, our study suggests *A. muciniphila*-derived OL may modulate inflammatory responses.

Remarkably, three host innate immunity genes — *Atf3, Tifa* and *Jmjd8* were co-regulated with *A. muciniplhila*. *Tifa* is located in the “cytokine-dependent colitis susceptibility locus” (*Cdcs1*) region, a critical genetic determinant of colitis susceptibility in 129 and B6 strains^60^. TIFA is an important modifier of innate immune signaling through its regulation of TRAF proteins, leading to the activation of NF-κB and inflammation. Considering the importance of TIFA-dependent immunity to Gram-negative bacteria^45^, and the differential effects of OL on LPS-treated BMDM from 129 and B6 strains, our results suggest that this gene could be a key player in *A. muciniphila*-OL-host interactions. Previous studies suggested that ATF3 modulates inflammatory responses by suppressing the expression of TLR4 or CCL4 in macrophages^44, 61^ and revealed a critical role of microbiota in ATF3-mediated gut homeostasis^62^. These studies showed that ATF3 negatively regulates *Il6* and *Il12* gene expression levels^44^. In line with this we found that OL negatively influence these cytokines in LPS-treated BMDM, and their abundance is associated with the same locus that influences *Atf3* expression. Although the molecular mechanisms underlying these associations warrant further investigation, these results suggest that *A. muciniphila* and OL levels are linked to these central players of the host immune defense system and support the predominant role of host genetics as a determinant of responses to gut microbes in particular to *A. muciniphila*.

In summary, the work presented here links the presence of OL in the human and mouse gut with *Akkermansia muciniphila* and suggest that these lipids are key players in *A. muciniphila*-host interactions. Our work highlights the importance of bacterial functions as mediators of host’s genetics influence on the gut microbiome.

## Methods

### Animals and sample collection

Animal care and study protocols were approved by the University of Wisconsin-Madison Animal Care and Use Committee. DO mice were obtained from the Jackson Laboratory (Bar Harbor, ME, USA) at ∼4 weeks of age and maintained in the Department of Biochemistry vivarium at the University of Wisconsin-Madison. Mice were housed on a 12-hour light:dark cycle under temperature- and humidity-controlled conditions. All mice were fed a high-fat high-sucrose diet (TD.08811, Envigo Teklad, 44.6% kcal fat, 34% carbohydrate, and 17.3% protein) ad libitum upon arrival to the facility. Mice were kept in the same vivarium room and were individually housed to monitor food intake and prevent cross-inoculation via coprophagy. DO mice were sacrificed at 22–25 weeks of age. Fecal samples were collected immediately before euthanasia after a 4 hour fast. Cecal contents, and additional tissues were harvested promptly after sacrifice and all samples were immediately flash frozen in liquid nitrogen and stored at −80°C until further processing. Other studies have been published with these mice^20, 63–65^.

#### Metagenomic shotgun DNA sequencing

Fecal DNA was extracted from individual fecal pellets collected from DO mice using previously described methods^7, 66^. Following DNA extraction, Illumina paired end libraries were constructed using a previously described protocol^67^, with a modification of gel selecting DNA fragments at ∼450 bp in length. Paired end (PE) reads (2 x 125) were generated using a combination of MiSeq and HiSeq 2500 platforms.

#### Metagenomic reads processing

Raw reads were preprocessed using Fastx Toolkit (ver. 0.0.13): (1) for demultiplexing raw samples, fastx_barcode_splitter.pl, with –partial 2, mismatch 2 was used; (2) when more than one forward and reverse read file existed for a single sample (due to being run on more than one lane, more than one platform, or at more than one time), read files were concatenated into one forward and one reverse read file; (3) barcodes were trimmed to form reads (fastx_trimmer -f 9 -Q 33); (4) and reads were trimmed to remove low quality sequences (fastq_quality_trimmer -t 20 -l 30 -Q33). Following trimming, unpaired reads were eliminated from the analysis using custom Python scripts. To identify and eliminate host sequences, reads were aligned against the mouse genome (mm10/GRCm38) using bowtie2^68^ (ver. 2.3.4) with default settings and microbial DNA reads that did not align to the mouse genome were identified using samtools (ver. 1.3) (samtools view -b -f 4 -f 8).

#### Metagenomic *de novo* assembly and gene prediction

After removing low quality sequences and host contaminating DNA sequences, each metagenomic sample was *de novo* assembled into longer DNA fragments (contigs) using metaSPAdes^69^ (ver. 3.11.1) with multiple k-mer sizes (metaspades.py -k 21, 33, 55, 77). Contigs shorter than 500bp were discarded from further processing. Open reading frames (ORFs) (i.e., microbial genes, also called metagenes), were predicted from assembled contigs via Prodigal^70^ (ver. 2.6.3) using Hidden Markov Model (HMM) with default parameters. All predicted genes shorter than 100bp were discarded from further processing. To remove redundant genes, all predicted ORFs were compared pairwise using criterion of 95% identity at the nucleotide level over 90% of the length of the shorter ORFs via CD-HIT^71^ (ver. 4.6.8). In each CD-HIT cluster, the longest ORF was selected as representative. This final non-redundant (NR) microbial gene set was defined as the DO gut microbiome NR gene catalog.

#### Metagenomic annotation

Gene taxonomic annotation was performed using DIAMOND^72^ (ver. 0.9.23) by aligning genes in the DO gut microbiome NR gene catalog to NCBI NR database (downloaded 2018-12-21) using default cutoffs: e-value < 1×10^-3^ and bit score > 50. Taxonomic assignment used the following parameters: “--taxonmap prot.accession2taxid.gz --taxonnodes nodes.dmp” in DIAMOND command and was determined by the LCA (Lowest Common Ancestor) algorithm when there were multiple alignments. Gene functional annotation was done using KEGG Orthology and Links Annotation (KOALA) method via the KEGG server (https://www.kegg.jp/ghostkoala/), using 2,698,820 prokaryotes genus pan-genomes as reference. The bit score cutoff for K-number assignment was 60.

#### Microbiome trait quantification

Quantification of microbial genes was done by aligning clean PE reads from each sample to the DO gut microbiome NR gene catalog using Bowtie2 (ver. 2.3.4) and default parameters. RSEM^73^ (ver. 1.3.1) was used to estimate microbial gene abundance. Relative abundance of microbial gene counts per million (CPM) were calculated using microbial gene expected counts divided by gene effective length then normalized by the total sum. We focused the taxonomic analysis on bacteria since these represented the vast majority of annotated metagenes. We detected 1,927,034 total metagenes, and from these 1,636,209 were annotated as bacterial genes, 195 archaeal genes, 17,372 eukaryotic genes, and 946 from viruses. There were also 272,312 genes that were unclassified. To obtain abundance information for microbial functions, CPM of genes with same KEGG Orthology (KO) annotation were summed together. In case there were multiple KO annotations for a single gene, we used all KO annotations. To obtain taxonomic abundance, CPM of genes with the same NCBI taxa annotation were summed together at phylum, order, class, family, and genus levels with a minimum of 10 genes in each taxon.

#### Metagenome Assembled Genomes (MAGs) reconstruction

To reconstruct bacterial genomes, we clustered assembled contigs using density-based algorithm DBSCAN using features of two reduced dimensions of contigs 5-mer frequency and contig coverage. The binning process was performed by the pipeline Autometa^74^ (docker image: ijmiller2/autometa:docker_patch) and allowed deconvolution of taxonomically distinct microbial genomes from metagenomic sequences. The quality of reconstructed metagenomes was evaluated using CheckM^75^ (ver. 1.1.3); genome completeness > 90% and genome contamination < 5%, were required to assign as high-quality MAGs. MAGs quantification was done by aligning all clean PE reads from each sample to MAGs from the same sample. Genome coverage was calculated using bedtools (v. 2.29.2) “genomecov” command followed normalization by library size across all samples. To further remove redundant MAGs, we clustered high-quality MAGs based on whole-genome nucleotide similarity estimation (pairwise ANIs) using Mash software^76^ (ver. 2.2) with 90% ANI. From high-quality MAGs, we also annotated predicted ORFs from each MAG against the KEGG database and compared the functional potential encoded among different taxa *A. muciniphila* MAG IDs are included in **Supplementary Table 14**.

#### Sample preparation for cecal lipidomic analysis

30 (± 7.5) mg cecal contents along with 10 μL SPLASH Lipidomix internal standard mixture were aliquoted into a tube with a metal bead and μL MeOH were added for protein precipitation. Contr 447 ol samples comprised 30 (± 7.5) mg of bead beat-combined DO founder strain cecum (NZO, PWK, NOD, B6, 129, AJ), extracted with each batch. To each tube, 900 μL methyl tert-butyl ether (MTBE) and 225 μL of water were added as extraction solvents. All steps were performed at 4 °C on ice. The mixture was homogenized by bead beating for 8 min at 25 Hz. Finally, the mixture was centrifuged for 8 min at 11,000 x g at 4 °C after which 240 μL of the lipophilic upper layer were transferred to glass vials and dried by vacuum centrifuge for 60 min.

The dried lipophilic extracts were re-suspended in 200 μL methanol (MeOH)/toluene (9:1, v/v) per 10 mg dry weight (minimum of 100 μL) to account for varying water content in the samples. The dry weight was determined by drying down the remaining mixture including all solid parts.

#### LC-MS/MS analysis of DO mouse cecal samples

Sample analysis by LC-MS/MS was performed in randomized order on an Acquity CSH C18 column held at 50 °C (2.1 mm x 100 mm x 1.7 μm particle diameter; Waters) using an Ultimate 3000 RSLC Binary Pump (400 μL/min flow rate; Thermo Scientific) or a Vanquish Binary Pump for validation experiments. Mobile phase A consisted of 10 mM ammonium acetate in ACN/H2O (70:30, v/v) containing 250 μL/L acetic acid. Mobile phase B consisted of 10 mM ammonium acetate in IPA/ACN (90:10, v/v) with the same additives. Mobile phase B was initially held at 2% for 2 min and then increased to 30% over 3 min. Mobile phase B was further increased to 50% over 1 min and 85% over 14 min and then raised to 95% over 1 min and held for 7 min. The column was re-equilibrated for 2 min before the next injection.

Twenty microliters of DO lipid extract were injected by an Ultimate 3000 RSLC autosampler (Thermo Scientific) coupled to a Q Exactive Focus mass spectrometer by a HESI II heated ESI source. Both source and inlet capillary were kept at 300 °C. Sheath gas was set to 25 units, auxiliary gas to 10 units, and the spray voltage was set to 5,000 V (+) and 4,000 V (-), respectively. The MS was operated in polarity switching mode acquiring positive and negative mode MS1 and MS2 spectra (Top2) during the same separation. MS acquisition parameters were 17,500 resolving power, 1 × 10^6^ automatic gain control (AGC) target for MS1 and 1 × 10^5^ AGC target for MS2 scans, 100-ms MS1 and 50-ms MS2 ion accumulation time, 200-to 1,600-Th MS1 and 200- to 2,000-Th MS2 scan range, 1-Th isolation width for fragmentation, stepped HCD collision energy (20, 30, 40 units), 1.0% under fill ratio, and 10-s dynamic exclusion.

#### Quantitative trait loci (QTL) mapping

Genetic QTL mapping was performed using the R/qtl2 (ver. 0.24) package^77^ which fit a linear mixed effect model that included accounted for overall genetic relationship with a random effect, i.e., kinship effect. The Leave One Chromosome Out (LOCO) method was used, which accounts for population structure without reducing QTL mapping power. For each gut microbiome trait, sex, days on diet, and mouse cohort (wave) were used as additive covariates. For cecum lipidome traits, sex and mouse cohort (wave) were used as additive covariates. For gut microbiome traits and cecum lipidome traits, normalized abundance/coverage was transformed to normal quantiles. The mapping statistic reported was the log10 likelihood ratio (LOD score). The QTL support interval was defined using the 95% Bayesian confidence interval^77^. Significance thresholds for QTL were determined by permutation analysis (n = 1000) and are reported at genome-wild adjusted *p* < 0.05. Suggestive QTL are reported at genome-wide adjusted *p* < 0.2.

#### Mediation analysis

Mediation analysis was carried out as previously described^78^. Mediation analysis was used to relate individual gut microbial metagenes and lipid features by scanning all 136,200 identified metagenes with at least 10 CPM in 20% of the samples to all 3,963 cecal lipid features. We used the subset of animals for which both gut metagenomic and cecal lipid data were available (n = 221). We first defined gut microbial traits with suggestive QTL as the outcome variable; we then included candidate cecal lipid mediators as additive covariates in the suggestive mbQTL mapping model and re-ran the QTL analysis. We performed the same analysis with cecal lipid features as the outcome and gut microbial features as candidate mediators. A mediatory role was supported by a significant decrease in LOD score from the original outcome QTL. Significance of the LOD score drop for a given candidate gut microbial metagene mediator on a given cecal lipid QTL was estimated by z-score scaled by LOD score drop and a conservative z-score ≤ −6 is recorded as a potential causal mediator. The mean of fitted distribution for a given gut bacterial taxon, e.g., all metagenes from gut *A. muciniphila*, is scaled to corresponding z-score to evaluate the mediation significance for this gut bacterial taxon.

#### Bacterial culturing and bacterial extracellular vesicles (EVs) isolation

*Akkermansia muciniphila* was grown anaerobically in defined medium (**Supplementary Table 15**). For test of effects of phosphate condition, the level was adjustedto 0.02, 0.2 or 2 mM. *E.coli* MS200-1 strain was grown in LC medium (10g/L Bacto-tryptone, 5g/L Bacto-yeast extract, 5g/L NaCl). *Bacteroides thetaiotaomicron* strain VPI-5482 was grown in CMM medium. All bacterial strains were grown in 37°C. Cells for lipid analyses from the three strains were obtained by centrifugation. Isolation of *A. muciniphila* extracellular vesicles using previously described method^79^.

#### Sample preparation for OL validation experiments

For cecal contents, 30 (± 6) mg cecal contents were aliquoted into a tube with a metal bead and 280 μL MeOH were added for protein precipitation. To each tube, 900 μL MTBE and 225 μL of water were added as extraction solvents. All steps were performed at 4 °C on ice. The mixture was homogenized by bead beating for 8 min at 25 Hz. For bacterial cultures, ∼ 75 ul of bacterial culture were aliquoted into a tube and 280 μL MeOH were added for protein precipitation. After the mixture was vortexed MTBE were added as extraction solvent and the mixture was vortexed for 10 s and mixed on an orbital shaker for 6 min. Phase separation was induced by adding 225 μL of water followed by 20 s of vortexing. All steps were performed at 4 °C on ice. Finally, each mixture was centrifuged for 8 min at 11,000 x g at 4 °C after which 240 μL of the lipophilic upper layer were transferred to glass vials and dried in a vacuum centrifuge for 60 min. The dried lipophilic extracts were re-suspended in 200 μL MeOH/toluene (9:1, v/v).

#### LC-MS/MS analysis of OL validation experiments

Sample analysis by LC-MS/MS was performed in randomized order on an Acquity CSH C18 column held at 50 °C (2.1 mm x 100 mm x 1.7 μm particle diameter; Waters) using an Ultimate 3000 RSLC Binary Pump (400 μL/min flow rate; Thermo Scientific) or a Vanquish Binary Pump. The same mobile phase and gradient as for the DO samples were used.

For the validation experiments, ten microliters of cecal or culture extract were injected by a Vanquish Split Sampler HT autosampler (Thermo Scientific) coupled to a Q Exactive HF mass spectrometer by a HESI II heated ESI source. Both source and inlet capillary were kept at 350 °C (Thermo Scientific). Sheath gas was set to 25 units, auxiliary gas to 15 units, and spare gas to 5 units, while the spray voltage was set to 3,500 V and S-lens RF level to 90. The MS was operated in polarity switching dd-MS2 mode (Top2) acquiring positive and negative mode MS1 and MS2 spectra during the same separation. MS acquisition parameters were 30,000 resolution and 1 × 10^6^ automatic gain control (AGC) target for MS1 and 5 × 10^5^ AGC target for MS2 scans, 100-ms MS1 and 50-ms MS2 ion accumulation time, 200 to 2,000-Th MS1 scan range, 1.0-Th isolation width for fragmentation, and stepped HCD collision energy (20, 30, 40 units).

#### Lipidomic analysis

All resulting LC-MS lipidomics raw files were converted to mgf files via MSConvertGUI (ProteoWizard, Dr. Parag Mallick, Stanford University) and processed using LipiDex^80^, Compound Discoverer 2.0 or 2.1.0.398 (Thermo Fisher Scientific) for DO and validation experiments, respectively. All raw files were loaded into Compound Discoverer with blanks marked as such to generate two result files using the following Workflow Processing Nodes: Input Files, Select Spectra, Align Retention Times, Detect Unknown Compounds, Group Unknown Compounds, Fill Gaps and Mark Background Compounds for the so called “Aligned” result and solely Input Files, Select Spectra, and Detect Unknown Compounds for an “Unaligned” Result. Under Select Spectra, the retention time limits were set between 0.4 and 21 min, MS order as well as unrecognized MS order replacements were set to MS1. Further replacements were set to FTMS Mass Analyzer and HCD Activation Type. Under Align Retention Times the mass tolerance was set to 10 ppm and the maximum shift according to the dataset to 0.6 min for the DO and 0.5 min for the validation experiments. Under Detect Unknown Compounds, the mass tolerance was also set to 10 ppm, with an S/N threshold of 5 (DO) or 3 (validation), and a minimum peak intensity of 5E6 (DO) or 1E5 (validation).

For the DO samples, [M+H]+1 and [M-H]-1 were selected as ions and a maximum peak width of 0.75 min as well as a minimum number of scans per peak equaling 7 were set. For the validation samples, [M+H]+1 and [M-H+TFA]-1 were selected as ions and a maximum peak width of 0.75 min as well as a minimum number of scans per peak equaling 5 were set. Lastly, for Group Unknown Compounds as well as Fill Gaps, mass tolerance was set to 10 ppm and retention time tolerance to 0.2 minutes. For best compound selection rules #1 and #2 were set to unspecified, while MS1 was selected for preferred MS order and [M+H]+1 as the preferred ion. For everything else, the default settings were used. Resulting peak tables were exported as excel files in three levels of Compounds, Compound per File and Features (just Features for the “Unaligned”) and later saved as csv. In LipiDex’ Spectrum Searcher “LipiDex_HCD_Acetate”, “LipiDex_HCD_Plants”, “LipiDex_Splash_ISTD_Acetate”, “LipiDex_HCD_ULCFA”, and “Ganglioside_20171205” were selected as libraries for the DO, and “Coon_Lab_HCD_Acetate_20171229”, “Ganglioside_20171205” and “Ornithine-Lipids_20180404” for the validation experiments. For all searches, the defaults of 0.01-Th for MS1 and MS2 search tolerances, a maximum of 1 returned search result, and an MS2 low mass cutoff of 61-Th were kept. Under the Peak Finder tab, Compound Discoverer was chosen as peak table type, and its “Aligned” and “Unaligned” results, as well as the MS/MS results from Spectrum Researcher uploaded. Features had to be identified in a minimum of 1 File while keeping the defaults of a minimum of 75% of lipid spectral purity, an MS2 search dot product of at least 500 and reverse dot product of at least 700, as well as a multiplier of 2.0 for FWHM window, a maximum 15 ppm mass difference, adduct/dimer and in-source fragment (and adduct and dimer) filtering, and a maximum RT M.A.D Factor of 3.5. As post-processing in the DO, all features that were only found in 1 file and had no ID were deleted, and duplicates deleted. Peak areas of the three targeted ornithine lipid species were obtained through TraceFinder v. 3.3.350.0 (Thermo Fisher Scientific). Details of the lipid classes searched for in these databases with their respective adducts were showed in **Supplementary Table 15**. Lipids ID matching was performed by < +/-5 ppm between runs.

#### RNA sequencing and eQTL analysis

Samples of flash-frozen distal ileum from DO mice were homogenized with Qiagen Tissuelyser (2 step 2min @ 25Hz, with flipping plate homogenization with 5 min ice incubation). Total RNA was extracted from homogenized samples using Qiagen 96 universal kit (Qiagen, Germantown, MD, USA). RNA clean up was performed by using Qiagen RNeasy Mini Kit (Qiagen). DNA was removed by on-column DNase digestion (Qiagen). Purified RNA was quantified using a Nanodrop 2000 spectrophotometer and RNA Fragment Analyzer (Agilent Technologies). Library preparation was performed by TruSeq Stranded mRNA Sample Preparation Guide (Illumina). IDT UDIs, Illumina UDIs or NEXTflex UDIs were used as barcodes for each library sample. RNA sequencing was performed on Illumina NovaSeq 6000 platform. Raw RNA Seq reads quality control was performed using Trimmomatic^81^ (ver. 0.39) with default parameters. Genotype-free genome reconstruction and allele specific expression (ASE) quantification were performed using the GBRS tool (http://churchill-lab.github.io/gbrs/). Genes with at least 10 transcripts/per million (TPM) in at least 10% of DO mice were used for downstream analyses. For eQTL mapping, sex, RNA-Seq index, RNA-Seq wave and mouse cohort (wave) were used as additive covariates. eQTL analysis was otherwise the same as previously described.

#### BMDM assay and cell viability measurement

Bone marrow was isolated from femur and tibia from ∼6 weeks-old B6 and 129 mice fed chow diet. Bone marrow cells were resuspended into single-cell suspension and cultured in complete DMEM medium supplemented with 10% FCS, 2mM l-glutamine, 1% penicillin/streptomycin, and 20 ng/ml mouse M-CSF (BioLegend) for the purpose of differentiation. BMDMs were harvested at day 7 and were treated with LPS, or OL or LPS+OL for 6 hours in media supplemented with 1% FBS, then supernatants were collected for measurement of cytokines. For optimization cytokines (TNF-α and IL6) production from LPS- or OL-treated BMDM were performed using Mouse TNF-α ELISA MAX Deluxe kit and Mouse IL-6 ELISA MAX Deluxe kit (BioLegend, San Diego, CA, USA). Follow-up cytokine (IL1β, IL6, IL10, IL-12, MCP-1, TNF-α, MIP-1α, GM-CSF, and RANTES) production assays in response to LPS+OL co-cultured BMDM were performed using Q-Plex™ Mouse Cytokine Screen 16-Plex (Quansys Biosciences, Logan, UT, USA). Cell viability was determined by flow cytometry (ThermoFisher Attune NxT) after staining with 7-amino-actinomycin D (eBioscience).

#### Data analysis and statistical analysis

All data integration and statistical analysis were performed in R (ver. 3.6.3). Differences between groups were evaluated using unpaired two-tailed Welch’s t-test. Enrichment analysis was performed using Fisher’s exact test using a custom R function. The correlation analysis was performed using Spearman’s correlation by R function “cor.test()”. For multiple testing, Benjamini-Hochberg FDR procedure was used to adjust p-values. Data integration was performed using R packages dplyr (ver. 1.0.6), tidyr (ver.1.1.3), reshape2 (ver. 1.4.4), data.table (ver. 1.14.0). Heatmap plots were performed using R package pheatmap (ver. 1.0.12). Other plots were performed using R packages ggplot2 (ver. 3.3.3), gridExtra (ver. 2.3), RcolorBrewer (ver. 1.1-2), ggsci (ver. 2.9).

### Data availability

DO metagenomic WGS data are available from the Sequence Read Archive (SRA) under accession PRJNA744213. RNA-Seq data are available from the Sequence Read Archive (SRA) under accession PRJNA772743. Mass spectrometry data files are available on Chorus (chorusproject.org) under accession with project ID 1681.

### Code availability

All code used in this study is available in GitHub (https://github.com/qijunz/Zhang_DO_paper) or on the corresponding software packages websites.

## Acknowledgements

The authors thank the University of Wisconsin Biotechnology Center DNA Sequencing Facility for providing sequencing and support services. The authors thank the University of Wisconsin Center for High Throughput Computing (CHTC) in the Department of Computer Sciences for providing computational resources, support, and assistance. We also thank the University of Wisconsin Carbone Cancer Center Flow Lab for support services.

## Funding

This work was supported by the National Institutes of Health (NIH) grants DK108259, HL144651 (F.E.R.), DK101573 (A.D.A), GM131817 (H.E.B.), GM070683 (K.W.B. and G.A.C.), NIH National Institute of Allergy and Infectious Diseases grant T32AI55397 (J.H.K.), NLM Computation and Informatics in Biology and Medicine Postdoctoral Fellowship 5T15LM007359 (L.L.T) and T32DK007665 (L.L.T.), and NIH Chemistry-Biology Interface Training Grant T32 GM008505 (T.J.P.).

## Author Contributions

F.E.R., M.P.K. and A.D.A conceived the study. Q.Z., V.L., L.L.T., A.D.A., J.J.C. and F.E.R. designed experiments. K.L.S., D.S.S., and M.E.R. assisted with mouse sample collection. L.L.T. and J.H.K. contributed to sample processing for DNA sequencing. Q.Z., L.L.T. and K.W.B. performed metagenomic and QTL analysis. V.L., K.A.O., E.A.T., T.R.R. and J.D.R. collected lipidomic data. V.L., I.J.M., M.P.K., D.M.G., G.R.K., D.T.P. and G.A.C. analyzed lipidome and lipidome QTL data. D.E.M., T.J.P., J.P.G. and H.E.B. synthetized OL. R.L.K. performed bacterial culture experiments. Q.Z. and K.K. performed cell culture studies. E.I.V. assisted with gnotobiotic mouse experiments. Q.Z., M.S. and A.J.L. assisted with intestine RNA-Seq. Q.Z., V.L., and F.E.R. wrote the manuscript. All authors approved the final manuscript.

## Competing interests

The authors declare no competing interests.

## Notes

### Competing Interest Statement

The authors have declared no competing interest.

