## Supplementary Information for "Systems genetics uncovers microbe-lipid-host connections in the murine gut"

### Supplementary Notes, Figures and Tables

#### Supplementary Note 1. DO metagenomic analysis

Shotgun DNA sequencing (Illumina 2X125 bp paired end reads) was performed in fecal samples collected from 297 DO mice at the end of the experiment. After removal of host DNA, each sample yielded on average 17.4 million high-quality microbial DNA paired end reads ($SD = 7.1 \times 10^6$). All shotgun sequencing reads were used to identify potential sample mix-ups and mislabeling during the library preparation process[1]. This revealed 15 samples containing mixtures of pairs of mice that were not used for further analyses. We also removed samples with low coverage (below 10 million). We proceeded with 264 DO metagenomic samples for QTL analysis. *De novo* metagenomic assembly captured 76% of the gut microbial DNA reads into contigs (**Supplementary Fig. 1a**) and included ~1.9 million unique predicted open reading frames (i.e., metagenes) with an average length of 690 bp. In order to assess the quality of the assembled non-redundant (i.e., NR) metagenes, we compared the percentage of sequence reads that map to our NR metagene assembly *vs*. a recently published mouse gut metagenome catalog[2]. Performing *de novo* assembly resulted in a larger fraction of the reads mapping to our assembly compared to the previously published catalog, i.e., alignment rate increased from 68.97% to 83.22% (**Supplementary Fig. 1b**) consistent with the notion that the DO mouse population contains different gut microbial communities relative to the previously published mouse cohort. Annotation of NR metagenes against the Kyoto Encyclopedia of Genes and Genomes (KEGG) orthology database via GhostKOALA identified 2803 functional orthologs (i.e., KOs) across all mice. It is important to note that each KO integrate abundance information of metagenes encoding functional orthologs – potentially from different species. Among these, Carbohydrate metabolism (14.30%), Membrane transport (9.05%) and Amino acid metabolism (8.43%) represented the categories with the largest number of functions detected in the distal gut of the DO mice (**Supplementary Fig. 1c**).

We used DIAMOND, a BLAST-based taxonomic classifier tool that uses the lowest common ancestor approach to assign phylogeny to all metagenes. First, we estimated the abundance of metagenomic reads in DO mice to each metagene and removed the lower abundance metagenes by the criteria that >10 counts per million in at least 10% of samples (**Supplementary Table 1**). We then summed the counts per million values of metagenes belonging to same taxon as the abundance of that taxon (Top 20 abundant genus showed in **Supplementary Fig. 1d**). We performed metagenomic binning to obtain species level bacterial genomes. This approach allowed us to detect diverse high-abundant bacterial genomes across all samples. After removing low quality bins, we obtained 5,611 putative metagenomic-assembled genomes (MAGs) representing genomes from 6 phyla including Actinobacteria, Bacteroidetes, Firmicutes, Proteobacteria, Tenericutes and Verrucomicrobia (**Supplementary Table 2**). Among these putative MAGs, 3,218 MAGs were identified as medium-quality MAGs with genome completeness above 50% and genome contamination below 5% as estimated by CheckM tool[3] (**Supplementary Fig. 1e**). 1,871 of these were high-quality with genome completeness above 90% and genome contamination below 5%. Furthermore, by reconstructing MAGs, we not only detect abundance of gut microbes but can also identify species-level variants. For example, by estimating pairwise average nucleotide identity of 46 high-quality *A. muciniphila* MAGs, we detected two distinct clusters (**Supplementary Fig. 1f**). These contain 3849 non-synonymous SNPs, indicating potential functional differences between these two *A. muciniphila* variants. Altogether, these results underscore the power of metagenomics for describing gut microbial communities, both taxonomically and in terms of their predicted gene functions. We also estimated narrow sense heritability of gut bacterial function traits via a linear mixed model and estimated heritability distribution of bacterial functions in each pathway (**Supplementary Table 3-4**).

**Supplementary Note 2. Correlation between cecal lipids and the gut microbiome**

To explore potential links between cecal lipid species and the gut microbiome, we performed correlation analysis between MAGs and cecal lipids abundance. Hierarchical clustering of correlation coefficients for each MAG-lipid species pair revealed groups of lipids associated with different gut microbiome profiles (**Supplementary Fig. 3**). MAGs clustered into five major groups while cecal lipids into six groups. More than 70% of the bacterial MAGs (**Supplementary Table 8**) were positively correlated with cluster 6 of cecal lipids. Fisher's exact test analysis disclosed that monogalactosyldiacylglycerols (MGDG) and phosphatidylglycerols (PG) were enriched in lipid cluster 6 ($P_{\text{BHadj}} = 1.47 \times 10^{-9}$ and $P_{\text{BHadj}} = 4.83 \times 10^{-6}$). These results are consistent with the notion that these lipids are highly prevalent across bacteria. In cecal lipid cluster 3, more than 90% of the lipids were phospholipids; plasmenyl-phosphatidylethanolamine ($P_{\text{BHadj}} = 3.92 \times 10^{-8}$) and phosphatidylcholines ($P_{\text{BHadj}} = 5.82 \times 10^{-8}$) were enriched in this cluster. The most highly positively correlated taxa with this lipid cluster were in MAG cluster 4, which contained 45 MAGs from the *Lachnospiraceae* family (67% of all MAGs in this cluster) including *Acetatifactor, Dorea, Lachnoclostridium, Roseburia*. In MAG cluster 1, 96% of the taxa were from the Bacteroidetes phylum, which were positively correlated with cecum lipid cluster 1, which were enriched in PC ($P_{\text{BHadj}} = 5.39 \times 10^{-7}$) and ceramide [BS] ($P_{\text{BHadj}} = 3.54 \times 10^{-3}$). Cecum lipid cluster 2 was enriched in fatty acids ($P_{\text{BHadj}} = 2.37 \times 10^{-25}$), ceramide [NS] ($P_{\text{BHadj}} = 2.30 \times 10^{-12}$) and ceramide [NP] ($P_{\text{BHadj}} = 5.75 \times 10^{-6}$) and was positively correlated with both Bacteroidetes and Firmicutes MAGs. Previous studies have shown that Bacteroidetes produce sphingolipids that are important for intestinal homeostasis and symbiosis[4]. To further confirm whether the significantly associated ceramides were derived from gut bacteria, we performed LCMS/MS from cecum in germ-free (GF) and conventionally-raised (convR) mice (**Supplementary Table 9**). We matched these lipid features to the ones detected in the DO mice. We found that over 67% of matched ceramides (present in DO and GF and/or convR) were significantly higher in convR mice compared to GF mice ($P_{\text{BHadj}} < 0.05$, **Supplementary Table 10**), indicating these associated cecal ceramides were derived from gut bacteria, most likely from Bacteroidetes. Altogether these results highlight potential taxa that modulate abundance of lipids in the gut.

**Supplementary Note 3. Cecal lipid features are associated with host genetics**

To link lipid features with the host genome, we performed QTL mapping of 3,384 mass spectral features. This resulted in 457 significant QTL for cecal lipid features (LOD > 7.5, $P_{Genome-wide-adj}$ < 0.05). In order to also capture the less direct associations of bacterial-derived lipids and the host genome, we included all suggestive QTL for further analysis (LOD > 6.0, $P_{Genome-wide-adj}$ < 0.2). This produced a total of 3,964 cecal lipid QTL (**Fig. 3c, Supplementary Table 11**). Notably, 68% of identified lipids gave a total of 1,162 QTL while a comparable proportion of 70% of unidentified features contributed 2,802 QTL. Some of the unidentified features QTL were in the region of QTL hotspots where identified lipid features co-mapped, suggesting features may share common drivers and/or belong to the same pathway and thus potentially enable the identification of unknown cecal lipid features, as previously reported[37]. Altogether these associations provide a wealth of information offering potential molecular descriptors of the genetic regulation of the microbiome.

**Supplementary Note 4. Ornithine lipids identification**

The lipid features whose QTL were mediated by gut *A. muciniphila* were initially not identified by our lipidomic analysis pipeline, they appeared to be closely related to each other. The three lipid features were observed closely in retention time and their precursor *m/z* in positive mode were 597.52, 611.54, and 625.55, consistently differing by 14 Da, the weight of a -$CH_2$ group. This led us to hypothesize that they were part of a fatty acid-based lipid class. When investigating their fragmentation spectra, we noticed recurring fragments at 70.065 and 115.087 Da, matching the formulas of $C_4H_8N$ and $C_5H_{11}N_2O$, respectively. We further identified characteristic neutral losses of water and one fatty acyl that also differed by 14 Da. Together, these findings suggest that the unidentified features could be ornithine lipids (OL). The three features would have the sum compositions of OL 30:0, OL 31:0, and OL 32:0, detected as [M+H]+ ions. In OL, a 3-hydroxy fatty acid is connected via an amide linkage to the ornithine amino acid that serves as the headgroup. A second fatty acid is then connected to the first via an ester linkage.

**Supplementary Note 5. Ornithine lipids synthesis**

**Chemical materials and methods.** All chemicals were obtained from Chem-Impex, Sigma-Aldrich, Agros Organics, or TCI America. All reagents and solvents were used without further purification except for hexane, ethyl acetate, and dichloromethane, which were distilled prior to use. Analytical thin-layer chromatography (TLC) was performed on 250 µm glass backed silica plates with F-254 fluorescent indicator from Silicycle. Visualization was performed using UV light and iodine.

**General instrumentation information.** NMR spectra were recorded in deuterated solvents at 400 MHz on a Bruker-Avance spectrometer equipped with a BFO probe, and at 500 MHz on a Bruker Avance spectrometer equipped with a DCH cryoprobe. Chemical shifts are reported in parts per million using residual solvent peaks or tetramethylsilane (TMS) as a reference. Couplings are reported in hertz (Hz). Electrospray ionization–exact mass measurement (ESI-EMM) mass spectrometry data were collected on a Waters LCT instrument.

**OL synthesis.** Tridecanoic acid (compound **1**, 3.2 g, 15 mmol) was dissolved in dichloromethane (150 mL, 0.1M) in a round bottom flask equipped with a stir bar. EDC-HCl (4.3 g, 22.5 mmol), DMAP (273 mg, 2.25 mmol), and Meldrum's acid (3.2 g, 22.5 mmol) were added to the flask, and the reaction was stirred overnight at room temperature. The next day, the reaction mixture was washed with 1M HCl (3 x 75 mL), saturated NaHCO3 (3x 75 mL), and brine (3 x 75 mL). The mixture was then dried over magnesium sulfate and concentrated under reduced pressure. The resultant oil was then dissolved in benzene (19 mL) in a round bottom flask with stir bar, and benzyl alcohol (45 mmol, 4.7 mL) was added. The reaction was heated to 95 °C for 3 hours, and then concentrated under reduced pressure. The crude reaction mixture was purified by silica gel flash chromatography (5-10% ethyl acetate in hexane as eluent), yielding 3.6 g of compound **2** as an oil (69% yield over two steps).

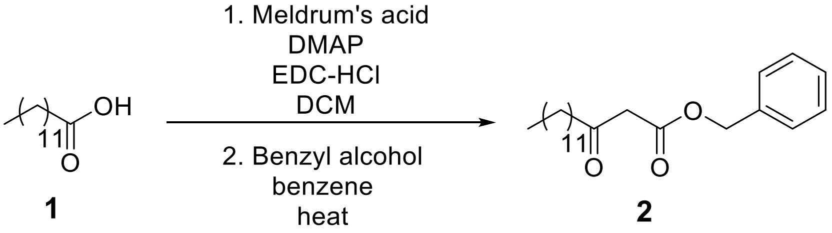


Compound **2** (3.6 g, 10.4 mmol) was added to a round bottom flask equipped with stir bar and dissolved in a 2:1 mixture of THF (16 mL) and ethanol (8 mL). The round bottom flask was cooled in an ice bath, and sodium cyanoborohydride (1.6 g, 26 mmol) was added to the mixture. 1M aqueous HCl (26 mL, 26 mmol) was added via addition funnel, and the reaction was allowed to stir to room temperature and monitored by TLC. Upon consumption of starting material, the aqueous portion of the reaction was extracted with dichloromethane (3 x 20 mL) and combined with the organic portion. The combined organic portions were washed with brine (3 x 20 mL), dried over MgSO4, and concentrated under reduced pressure to yield 3.26 g of compound **3** (93% crude). The material was used without further purification.

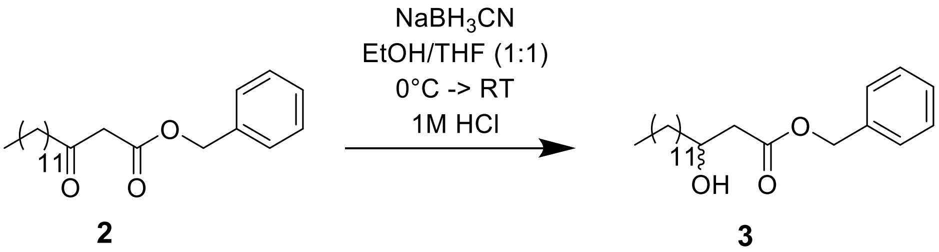


Pentadecanoic acid (1.93g, 9 mmol) was added to a round bottom flask equipped with a stir bar and dissolved in dichloromethane (80 mL). To the flask was added EDC-HCl (2.68 g, 14 mmol), DMAP (974 mg, 8 mmol), and compound **3** (2.78 g, 8 mmol). The reaction mixture was allowed to stir overnight at room temperature. The next day, the mixture was washed with 1M HCl (3 x 50 mL), saturated NaHCO3 (3 x 50 mL), and saturated brine (3 x 50 mL). The mixture was then dried over magnesium sulfate and concentrated under reduced pressure. The crude material was purified by silica gel flash chromatography (5-10% ethyl acetate in hexane as eluent), yielding 4.3 g of compound **4** (94% isolated yield).

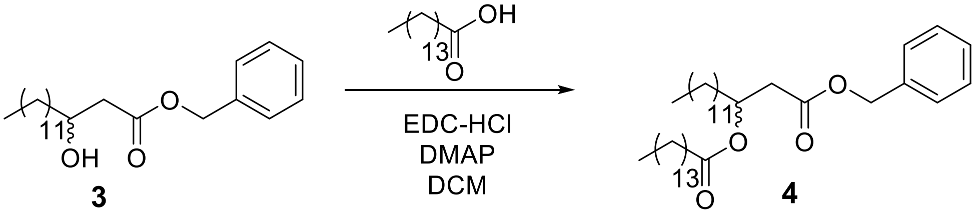


To a flame dried round bottom flask equipped with a stir bar was added Pd/C (798 mg, 0.75 mmol Pd). Dry dichloromethane was added to the flask to make a slurry, and the atmosphere was exchanged for nitrogen. Compound **4** (4.3 g, 7.5 mmol) was dissolved in anhydrous methanol and added to the reaction vessel. The atmosphere was then exchanged for hydrogen (balloon pressure), and the reaction was allowed to proceed overnight. The next day, the reaction was diluted with ethyl acetate and filtered over celite. The mixture was concentrated under reduced pressure to yield compound **5** as a white solid (3.5 g, 97% crude yield). The material was used without further purification.

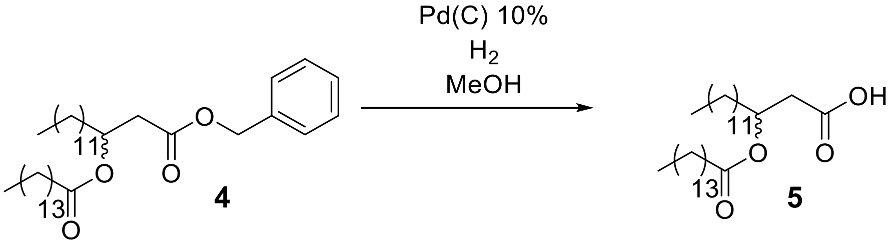


Compound **5** (256 mg 0.5 mmol) was added to a round bottom flask equipped with a stir bar and dissolved in DMF (5 mL). To the flask was added DIPEA (277 μL, 1.6 mmol) and HATU (216 mg, 5.5 mmol), and the mixture was stirred for 15 minutes. Protected ornithine (250 mg, 0.6 mmol) was added to the mixture, which was stirred at room temperature and monitored by TLC. When starting material was no longer observed by TLC, the mixture was diluted in diethyl ether (20 mL) and washed with 1M HCL (3 x 20 ml), saturated NaHCO3 (3 x 20 mL), and brine (3 x 20 mL). The mixture was dried over magnesium sulfate and concentrated under reduced pressure to yield a white solid (376 mg crude). This sample was combined with an additional sample of the same crude material that appeared identical by $^{1}$H NMR analysis and was then purified by silica gel flash chromatography (25% ethyl acetate in hexanes as eluent) to yield 131 mg of compound **6**.

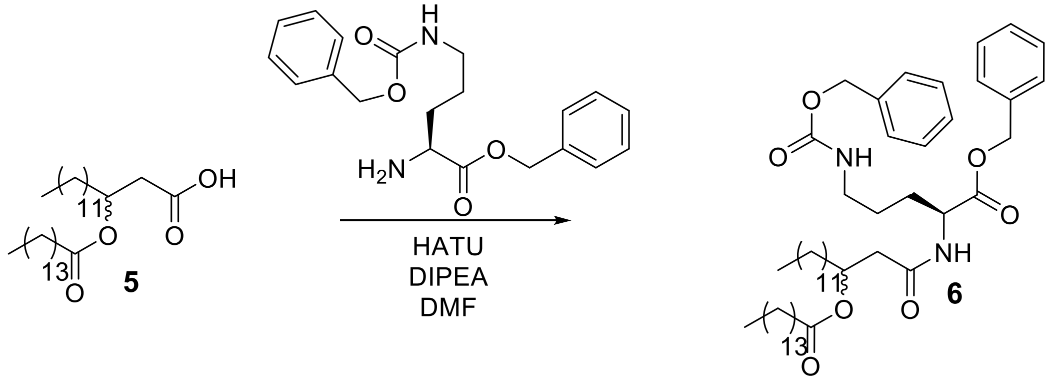


To a flame dried round bottom flask equipped with a stir bar was added Pd/Cn (17.0 mg, 0.16 mmol Pd). Dry dichloromethane was added to the flask to make a slurry, and the atmosphere was exchanged for nitrogen. The protected ornithine lipid (compound **6**, 131 mg, 0.160 mmol) was dissolved in a mixture of 4 mL anhydrous methanol/DCM (1:1) and added to the reaction vessel. The atmosphere was then exchanged for hydrogen (balloon pressure), and the reaction was allowed to proceed overnight. The next day, the reaction was filtered over celite. The mixture was concentrated under reduced pressure to yield **OL** as an off-white solid (82.2 mg, 86% crude yield). Deprotected **OL** was identified using LC and ESI-EMM ([M]+ calculated 597.5207, measured 597.5188, 0.002 ppm) in the resultant mixture and the material was used without further purification in the experiments described herein.

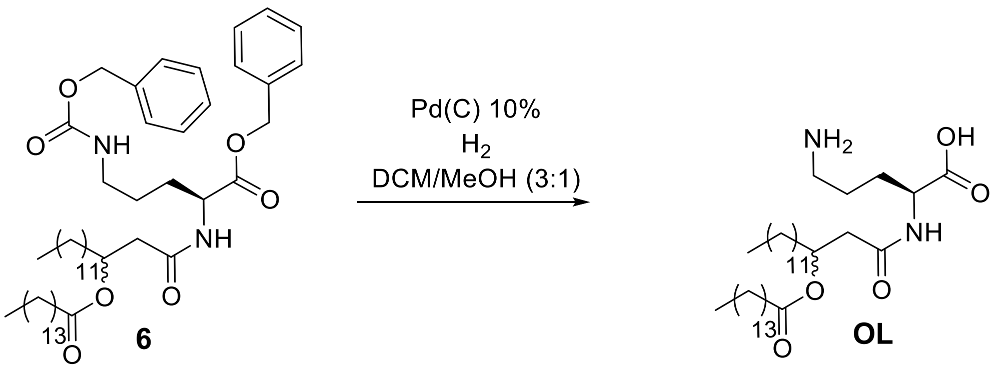


**Compound characterization data**

Compound **2**

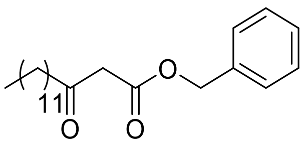

$^1$H NMR (500 MHz, Chloroform-d) δ 7.39 – 7.31 (m, 5H), 5.17 (s, 2H), 3.48 (s, 2H), 2.49 (t, J = 7.4 Hz, 2H), 1.61 – 1.52 (m, 2H), 1.25 (d, J = 5.0 Hz, 18H), 0.88 (t, J = 6.9 Hz, 3H). $^{13}$C NMR (126 MHz, $CDCl_3$) δ 202.71, 167.10, 135.36, 129.04, 128.62, 128.46, 128.39, 128.23, 128.17, 125.30, 88.81, 67.10, 65.67, 53.42, 49.25, 43.10, 35.11, 31.92, 30.93, 29.65, 29.63, 29.60, 29.44, 29.35, 29.07, 29.01, 26.25, 23.46, 22.70, 14.12

Compound **3**

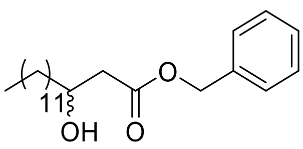

$^1$H NMR (400 MHz, Chloroform-d) δ 7.41 – 7.30 (m, 5H), 5.16 (s, 2H), 4.02 (tq, J = 7.7, 3.6 Hz, 1H), 2.89 (s, 1H), 2.62 – 2.40 (m, 2H), 1.59 – 1.37 (m, 3H), 1.37 – 1.19 (m, 18H), 0.88 (t, J = 6.7 Hz, 3H). $^{13}$C NMR (126 MHz, $CDCl_3$) δ 173.02, 135.58, 128.65, 128.42, 128.28, 68.12, 67.99,

66.57, 53.43, 41.33, 36.57, 31.93, 29.68, 29.66, 29.59, 29.57, 29.52, 29.36, 25.62, 25.47, 22.70, 14.13.

Compound **4**

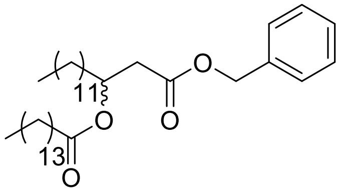

[1]H NMR (500 MHz, Chloroform-d) δ 7.38 – 7.29 (m, 5H), 5.23 (tt, J = 7.5, 5.4 Hz, 1H), 5.11 (s, 2H), 2.67 – 2.53 (m, 2H), 2.29 – 2.11 (m, 2H), 1.66 – 1.49 (m, 5H), 1.41 – 1.19 (m, 44H), 0.88 (t, J = 6.8 Hz, 6H). [13]C NMR (126 MHz, $CDCl_3$) δ 173.15, 170.31, 135.79, 128.55, 128.34, 128.28, 70.28, 66.44, 39.35, 34.45, 34.06, 31.93, 29.71, 29.70, 29.68, 29.67, 29.66, 29.64, 29.55, 29.51, 29.48, 29.37, 29.36, 29.30, 29.14, 25.12, 24.99, 22.70, 22.66, 14.12. ESI-EMM: $[M+NH_4]+$ calculated 590.5139; measured 590.5139, 0.7 ppm.

Compound **5**

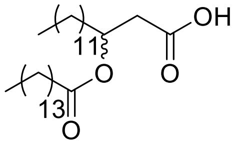

[1]H NMR (500 MHz, Chloroform-d) δ 5.21 (p, J = 6.2 Hz, 1H), 2.60 (qd, J = 15.7, 6.3 Hz, 2H), 2.28 (t, J = 7.5 Hz, 2H), 1.61 (q, J = 7.3 Hz, 4H), 1.25 (s, 44H), 0.88 (t, J = 6.8 Hz, 6H). [13]C NMR (126 MHz, $CDCl_3$) δ 173.34, 70.11, 38.84, 34.50, 34.04, 31.94, 29.72, 29.71, 29.69, 29.67, 29.65, 29.56, 29.51, 29.37, 29.30, 29.15, 25.14, 25.02, 22.70, 14.12, 0.00. ESI-EMM: [M-H]- calculated 481.4262; measured 481.4297, 0.2 ppm.

Compound **6**

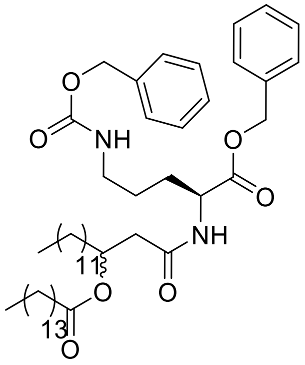


$^{1}$H NMR (500 MHz, Chloroform-d) δ 7.38 – 7.27 (m, 4H), 5.25 – 5.09 (m, 10H), 5.07 (s, 2H), 3.16 (q, J = 6.7 Hz, 2H), 2.50 (dq, J = 13.6, 6.9, 6.3 Hz, 2H), 2.27 (td, J = 7.4, 2.0 Hz, 2H), 1.87 (dp, J = 14.5, 4.9, 4.5 Hz, 1H), 1.74 – 1.63 (m, 1H), 1.59 (ddq, J = 17.2, 12.1, 6.0, 5.1 Hz, 4H), 1.26 (q, J = 5.5, 3.2 Hz, 44H), 0.88 (t, J = 6.9 Hz, 6H). 13C NMR (126 MHz, $CDCl_3$) δ $^{13}$C NMR (126 MHz, $CDCl_3$) δ 174.35, 173.60, 173.49, 173.28, 173.27, 172.33, 172.07, 172.04, 171.19, 169.78, 169.74, 156.55, 146.18, 136.60, 135.22, 135.20, 128.65, 128.54, 128.50, 128.32, 128.08, 128.06, 71.14, 70.18, 67.28, 67.26, 66.64, 60.41, 51.89, 41.67, 41.45, 40.38, 38.94, 34.52, 34.49, 34.33, 34.14, 34.04, 31.94, 29.72, 29.71, 29.70, 29.67, 29.58, 29.53, 29.52,z 29.37, 29.33, 29.31, 29.19, 29.18, 29.15, 28.89, 28.22, 25.94, 25.88, 25.81, 25.59, 25.25, 25.22, 25.14, 25.04, 25.01, 24.99, 22.70, 14.12. ESI-EMM: [M+Na]+ calculated 843.5858; measured 843.5851, 0.8 ppm.

**Supplementary Note 6. Ornithine lipids are detected in *A. muciniphila*-derived Extracellular Vesicles (AmEVs).**

In Gram-negative bacteria, extracellular vesicles (EVs) are derived from the outer membrane. Previous studies suggest that bacterial EVs can penetrate the mucus barrier, get internalized in the epithelium and can also access immune cells in the lamina propria as well as play a crucial role in immunity and maintenance of gut homeostasis [5–7]. *A. muciniphila* produces extracellular vesicles that have beneficial effects on the host[8]. Animal studies showed that *A. muciniphila*-derived EVs are able to improve the intestinal barrier by increasing tight junctions[9] and ameliorating inflammation caused by colitis[10]. We examined whether OL were present in *A. muciniphila*-derived EVs. We isolated EVs from this bacterium grown in vitro and measured the lipid features using LC-MS/MS. We found AmEVs to contain OL at levels comparable to whole *A. muciniphila* cells (**Supplementary Fig. 4b**). Ornithine lipids were among the most abundant lipid features detected in *A. muciniphila*-derived EVs with several features significantly enriched including OL_30:0, OL_31:0, and OL_32:0 which were also detected in bacterial culture, and mouse colonization studies. These results suggested that OL are likely localized in *A. muciniphila* outer membrane and provide insights into how these lipids may access the host.

**Supplementary Note 7. Co-mapping QTL highlights**

**Co-mapping of intestinal expression QTL (eQTL) and microbiome QTL (mbQTL).** One interesting example of co-mapping was observed between bacterial lipopolysaccharide cholinephosphotransferase (K07271, LicD) and host peptidoglycan recognition protein 1 (PGLYRP1) gene on Chr8. K07271 has a QTL with LOD 7.9 and a QTL peak at 12.3 Mbp on Chr8 whereas the PGLYRP1 gene has a trans-eQTL on chr8 with LOD of 6.3 at 14.1 Mbp (**Supplementary Fig. 6a**). Notably, these co-mapping traits also share the same allele effect pattern in which 129 and NOD haplotypes have strong positive and negative associations, respectively. Correlation of QTL allele effects between these two traits was significant ($R$ = 0.96). Bacterial lipopolysaccharide cholinephosphotransferase can incorporate environmental choline into lipopolysaccharide (LPS) as phosphorylcholine (ChoP) whereas LicD affects bacterial surface structure by linking ChoP to LPS, which can in turn change susceptibility of the organism to host defense mechanisms[11]. Hosts' peptidoglycan recognition proteins (PGRPs) bind to both Gram-positive and Gram-negative bacteria and have bactericidal activity[12]. PGRPs have one or two PGRP domains with a binding site specific for bacterial peptidoglycan[13]. Furthermore, a recent study showed that *Pglyrp1*$^{-/-}$ mice have alterations in the gut microbiome and exhibit decreased responsiveness to allergic asthma[14]. While it is still not clear how changes in Pglyrp1 gene expression impact the gut microbiome or inversely, our results suggest that differences in Pglyrp1 gene expression are significantly associated with bacteria encoding LicD. Interestingly, we also detected several bacterial motility related genes including *fliC* (K02406), *fliG* (K02410), *flgE* (K02390), *flhA* (K02400), *fliF* (K02409) that showed opposite QTL allele effect in this region, with the 129 allele negatively associated with abundance of these traits (**Supplementary Fig. 6a**). This suggests that gut bacterial motility functions are negatively associated with host *Pglyrp1* gene expression level. Notably, two MAGs also mapped to this locus with the same allele effects as these motility related genes. These were annotated at the genus level as *Acetatifactor*. *Acetatifactor muris* was first isolated from the intestine of an obese mouse[15]. Recent studies showed that the abundance of *Acetatifactor muris* in the gut is influenced by dietary fat sources[16] and correlated with bile acids including lithocholic acid (LCA) and ursodeoxycholic acid (UDCA)[17].

**Co-mapping of intestinal eQTL with cecal liplid QTL (clQTL).** There were several instances in which unidentified lipid features co-mapped with other traits. Below we provide two examples: Example 1: At Chr4: ~50Mbp, we detected a strong CAST allele effect on an unidentified feature with a retention time of 0.53 minutes and an m/z of 414.19421 in positive polarity that is only shared by few other traits. Notably, among them is the local eQTL for *Acnat1*, acyl-coenzyme A amino acid N-acyltransferase 1 (**Supplementary Fig. 6b**). ACNAT1 catalyzes the addition of taurine to fatty acids to form N-acyl taurines (NAT)[18] (**Supplementary Fig. 6c**). To a lesser extent, it can also conjugate bile acids. Given this lead, the fragmentation spectrum of the feature allowed an identification as NAT 18:5;O2 (**Supplementary Fig. 6d**). Particularly intriguing was the co-mapping of microbial K03704 (Cspa, cold shock protein). CspA acts as an RNA chaperone that has been linked to the regulation of growth and stress adaptation as well as to the promotion of survival during stationary-phase. Cold shock proteins are also important for virulence in macrophages and mice. While the nature of these associations remains unknown, these results suggest that cold shock protein may be involved in a stress response to the taurine conjugated metabolite.

Example 2: Another instance of co-mapping occurred on Chr17: ~32Mbp and included several unidentified lipid QTL (**Supplementary Fig. 6e**). We used the allele effect pattern of co-mapping traits to identify subgroups of lipid features and genes expressed in the gut having the same genetic architecture. This analysis revealed an unidentified lipid feature with an m/z of 605.41 that showed the highest significance with several genes in this region including Cyp4f, two genes encoding for P450 enzymes that show matching or inverse allele effects (Cyp4f13, Cyp4f14)[19]. Importantly, we were able to identify several of the unidentified features as tocopherol species, and the above-mentioned feature as alpha-tocopherol glucuronide (**Supplementary Fig. 6f**). This observation is in line with vitamin E metabolites being excreted as sulfate and glucuronide conjugates[20]. Furthermore, disruption of the Cyp4f14 gene in mouse causes severe perturbations in vitamin E metabolism. In summary, we were able to use a combination of genomic and mass spectral data to annotate modified cecal lipid species that were unidentified after a standard database search. In several instances, co-mapping microbial QTL raised the question of an involvement of the gut microbiome in their metabolic processes.

**Supplementary Figure 1. DO metagenomic analysis. a,** Average percent of assembled reads across all samples. **b,** Comparison of percent of reads mapping to our generated assembly *vs.* public database. **c**, Microbial functions detected for KEGG pathways across all metagenomes. KEGG Orthology (KO) numbers were identified by annotating predicted ORFs to the KEGG database. **d,** Top 20 gut microbial genera detected across all DO mice. **e,** Quality of metagenome-assembled genomes. **f,** Two variants of *A. muciniphila* MAGs detected in the DO mice.

**Supplementary Figure 2. DO gut microbiome QTL hotspot at Chr15: ~63Mbp. a,** Founder allele effects of KO and taxa trait QTL at Chr15 hotspot (LOD > 6, $P_{\text{Genome-wide-adj}} < 0.2$). **b,**

Abundance of KOs that mapped to Chr15 hotspot across all MAGs. Sporulation functions were not detected in Bacteroidetes. **c,** Estimated founder allele effects for Bacteroidetes and Firmicutes, and Bacteroidetes/Firmicutes ratio (left panel). Observed abundance of Bacteroidetes Firmicutes and Bacteroidetes/Firmicutes ratio in founder strains as determined by Kemis et al (right panel). **d,** SNPs significantly associated with these traits in Chr15 hotspot include two intron SNPs in *Gsdmc* and *Gsdmc2* genes.

**Supplementary Figure 3. Correlation between gut MAGs and cecum lipids. a,** Heatmap showing Spearman correlation coefficients between the abundance of MAGs and cecal lipid levels across DO mice. Bacterial MAGs were clustered into 5 groups whereas cecal lipids were clustered into 6 groups. **b,** Enrichment of the lipid classes for each cecal lipid clusters. Fisher's exact test was used and Benjamini-Hochberg for multiple tests correction.

**Supplementary Figure 4*. *Detection of ornithine lipids (OL) in *Akkermansia muciniphila.***
**a,** Heatmap showing relative abundance of all OL species detected in cell pellets from *A. muciniphila* grown *in vitro* in defined media supplemented with different levels of phosphate: 20μM, 200μM and 2000μM. **b,** Relative abundance of lipid features detected in cell pellets from *A. muciniphila* grown in defined media with different levels of phosphate. Top 200 most abundant lipids features are shown. **c,** Relative abundance of OL features detected extracellular vesicles (AmEVs) purified from *A. muciniphila* grown in defined medium with the comparison to *A. muciniphila* cells.

**Supplementary Figure 5. Cytokine production by bone-marrow-derived macrophages (BMDM). (a)** TNF-$\alpha$ and **(b)** IL-6 levels detected in supernatants from BMDM cells in B6 and 129 mice treated for 6 hours with different concentrations of LPS or OL. **c,** Cell viability of BMDM cells in B6 and 129 mice treated for 6 hours with 10 ng/mL LPS and different concentrations of OL.

**Supplementary Figure 6. Examples of co-mapping QTL.** **a**, At Chr8: 10.5-14.5 Mbp, co-mapping of gut bacterial lipopolysaccharide cholinephosphotransferase function with *Pglyrp1* eQTL was observed. **b**, At Chr4: 50 Mbp, co-mapping of an unidentified cecal feature and a local *Acnat1* eQTL was observed. **c**, The knowledge of *Acnat1* conjugating taurine to fatty acids guided the identification of the feature as an N-acyl taurine. **d**, Fragmentation pattern of identified N-acyl taurine. **e**, At Chr17: 30-34 Mbp, several unidentified features co-mapped which subsequently could be identified as tocopherols and exemplarily shown for the most significant feature alpha-tocopherol glucuronide. **f**, Fragmentation pattern of identified alpha-tocopherol glucuronide.

**Supplementary Figure 7. Founder allele effects on co-mapping traits associated with *A. muciniphila* levels.** *A. muciniphila*, cecum OLs, and eQTL genes co-mapping at Chr1: 90-95 Mbp, Chr2: 77-81 Mbp, Chr7: 126-131 Mbp, Chr12: 55-63 Mbp, and Chr15: 75-79 Mbp.

**Supplementary Table 1. DO gut microbiome metagene annotation.**

**Supplementary Table 2. DO gut microbiome MAG annotation.**

**Supplementary Table 3. Narrow sense heritability for bacterial functions detected in the gut of DO mice consuming a High Fat/High Sucrose diet (n=264).**

**Supplementary Table 4. Narrow sense heritability for bacterial taxa detected in the gut of DO mice consuming a High Fat/High Sucrose diet (n=264).**

**Supplementary Table 5. QTL for microbial functions detected in the gut of DO mice consuming a High Fat/High Sucrose diet (n=264).**

**Supplementary Table 6. QTL for microbial taxa detected in the gut of DO mice consuming a High Fat/High Sucrose diet (n=264).**

**Supplementary Table 7. QTL hotspots identified for gut microbial features.**

**Supplementary Table 8. Correlation between Metagenome-Assembled Genomes (MAGs) clusters and lipid clusters.**

**Supplementary Table 9. Lipids detected in cecal contents from germ-free (GF) and conventionally-raised (ConvR) mice consuming High Fat/High Sucrose diet.**

**Supplementary Table 10. Cecal ceramides detected in DO mice are largely of microbial origin.**

**Supplementary Table 11. QTL for lipid features detected in the gut of DO mice consuming a High Fat/High Sucrose diet (n=381).**

**Supplementary Table 12. eQTL for small intestine genes of DO mice consuming a High Fat/High Sucrose diet (n=234).**

**Supplementary Table 13. Correlation of small intestine eQTL allele effects with *A. muciniphila*/OL QTL allele effects.**

**Supplementary Table 14. List of *A. muncinphila* MAGs.**

**Supplementary Table 15. Media recipe for bacterial culture.**

**Supplementary Table 16. Lipid class abbreviations.**
