## Supplementary Figures for "Systems genetics uncovers microbe-lipid-host connections in the murine gut"

Supplementary Figure 1

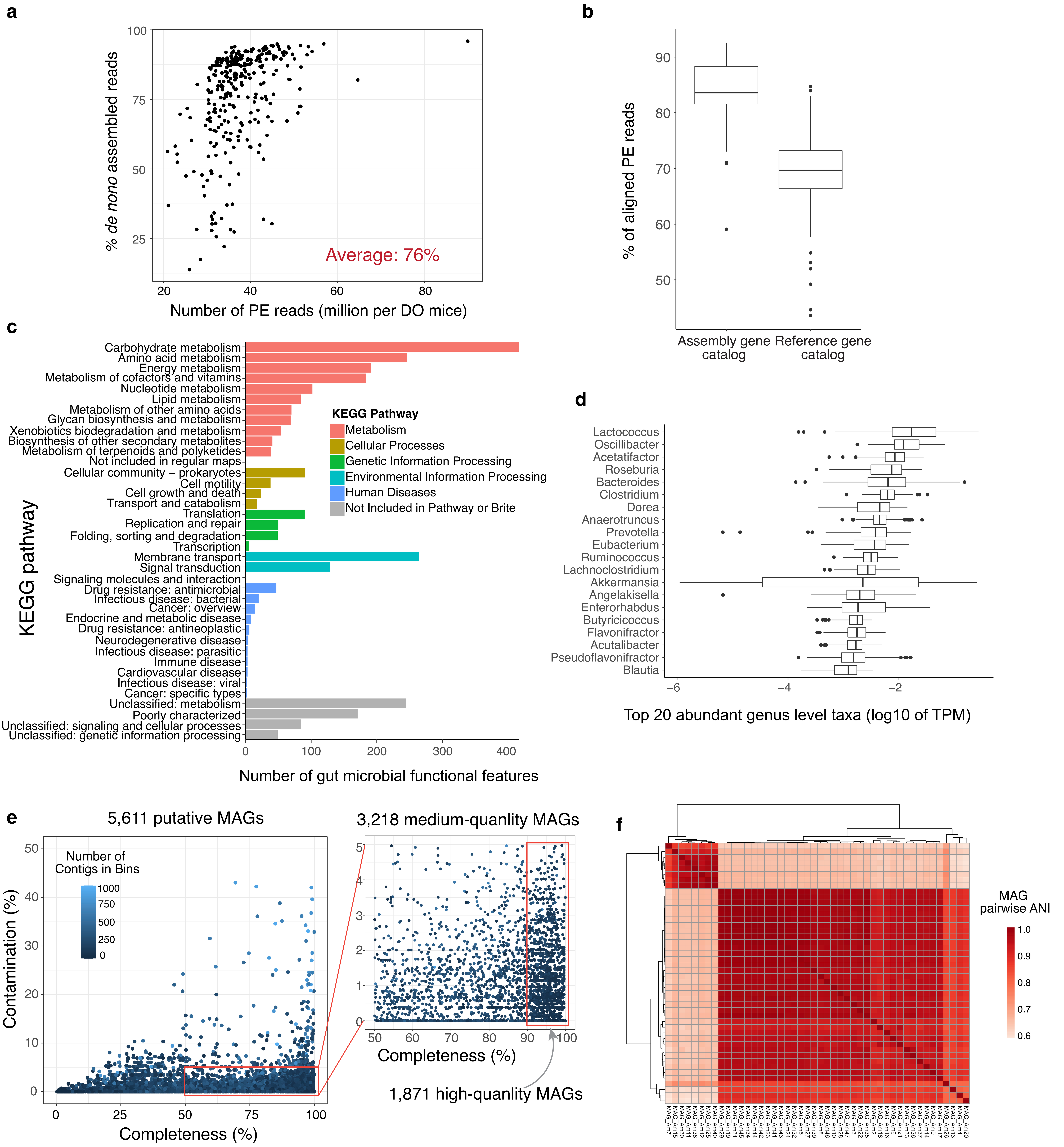

Supplementary Figure 2

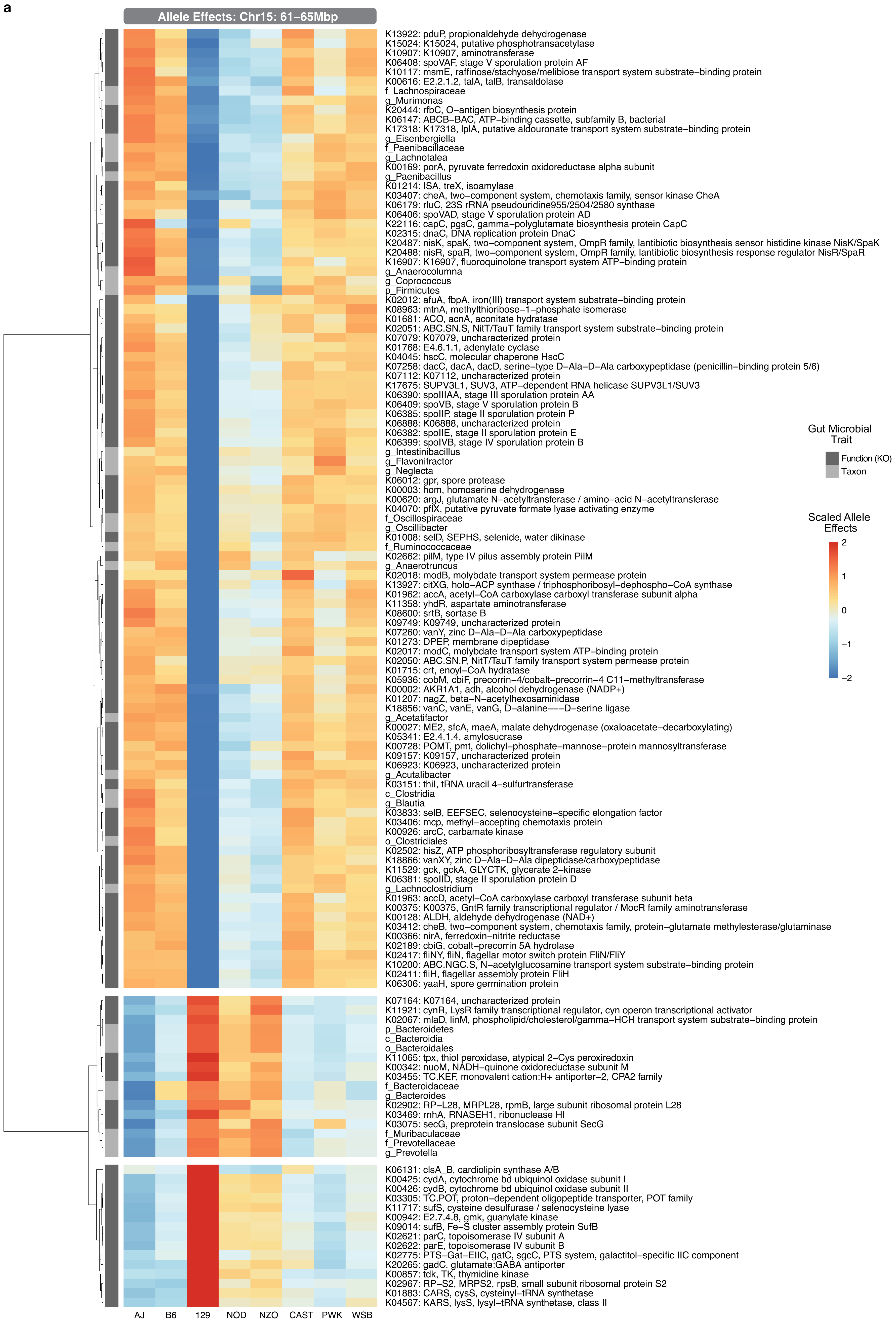

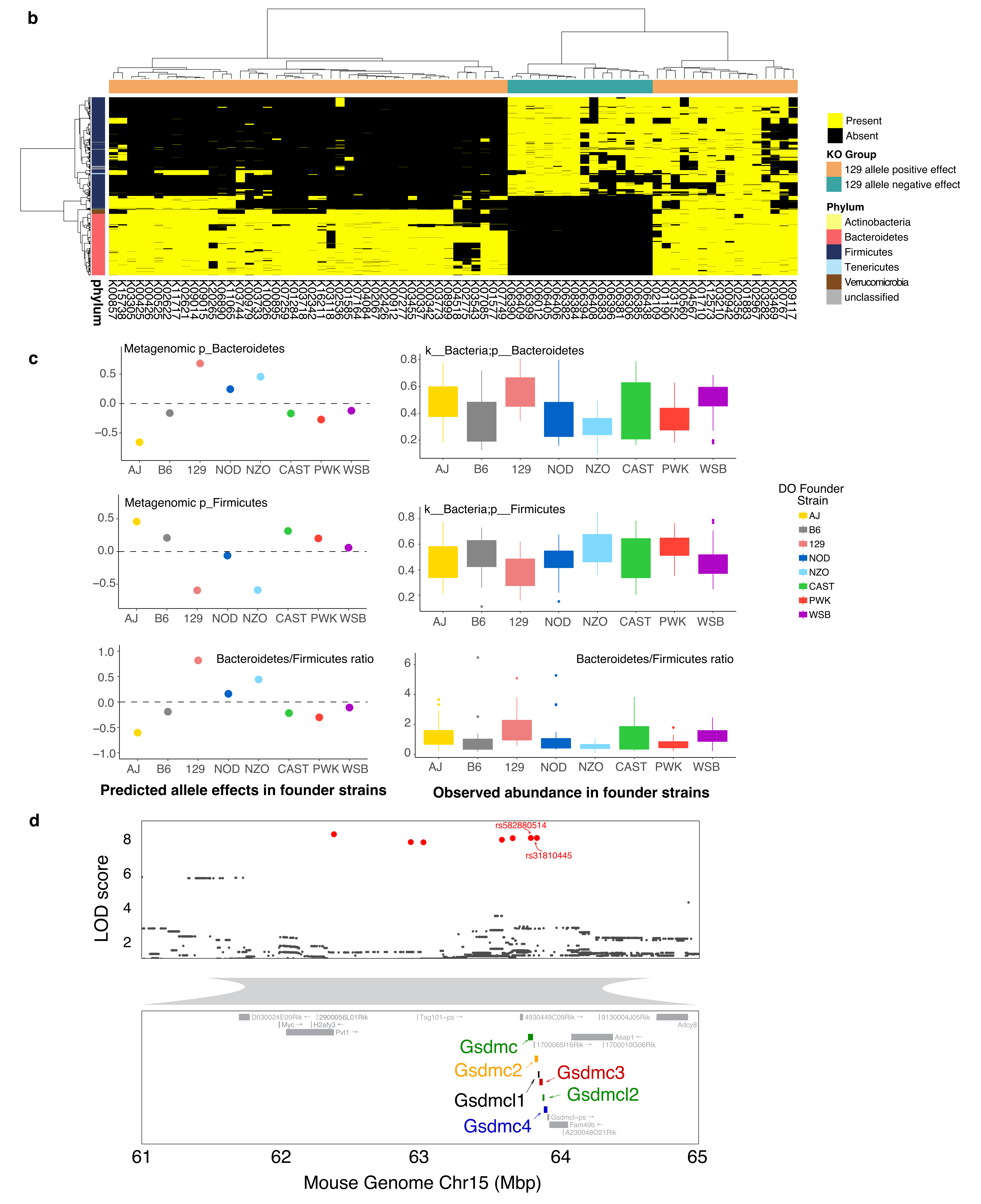
b
Present
Absent
KO Group
129 allele positive effect
129 allele negative effect
Phylum
Actinobacteria
Bacteroidetes
Firmicutes
Tenericutes
Verrucomicrobia
unclassified
phylum
c
Metagenomic p_Bacteroidetes
k__Bacteria;p__Bacteroidetes
Metagenomic p_Firmicutes
k__Bacteria;p__Firmicutes
Bacteroidetes/Firmicutes ratio
AJ B6 129 NOD NZO CAST PWK WSB
DO Founder Strain
AJ
B6
129
NOD
NZO
CAST
PWK
WSB
Predicted allele effects in founder strains
Observed abundance in founder strains
d
rs582880514
rs31810445
LOD score
D030024E09Rik
2900056L01Rik
Myc
H2afy3
Pvt1
Tsg101-ps
4930449C09Rik
9130004J05Rik
Adcy8
Asap1
1700065I16Rik
1700010G06Rik
Gsdmc
Gsdmc2
Gsdmc3
Gsdmcl1
Gsdmcl2
Gsdmc4
Gsdmcl-ps
Fam49b
A230048O21Rik
61
62
63
64
65
Mouse Genome Chr15 (Mbp)

Supplementary Figure 3

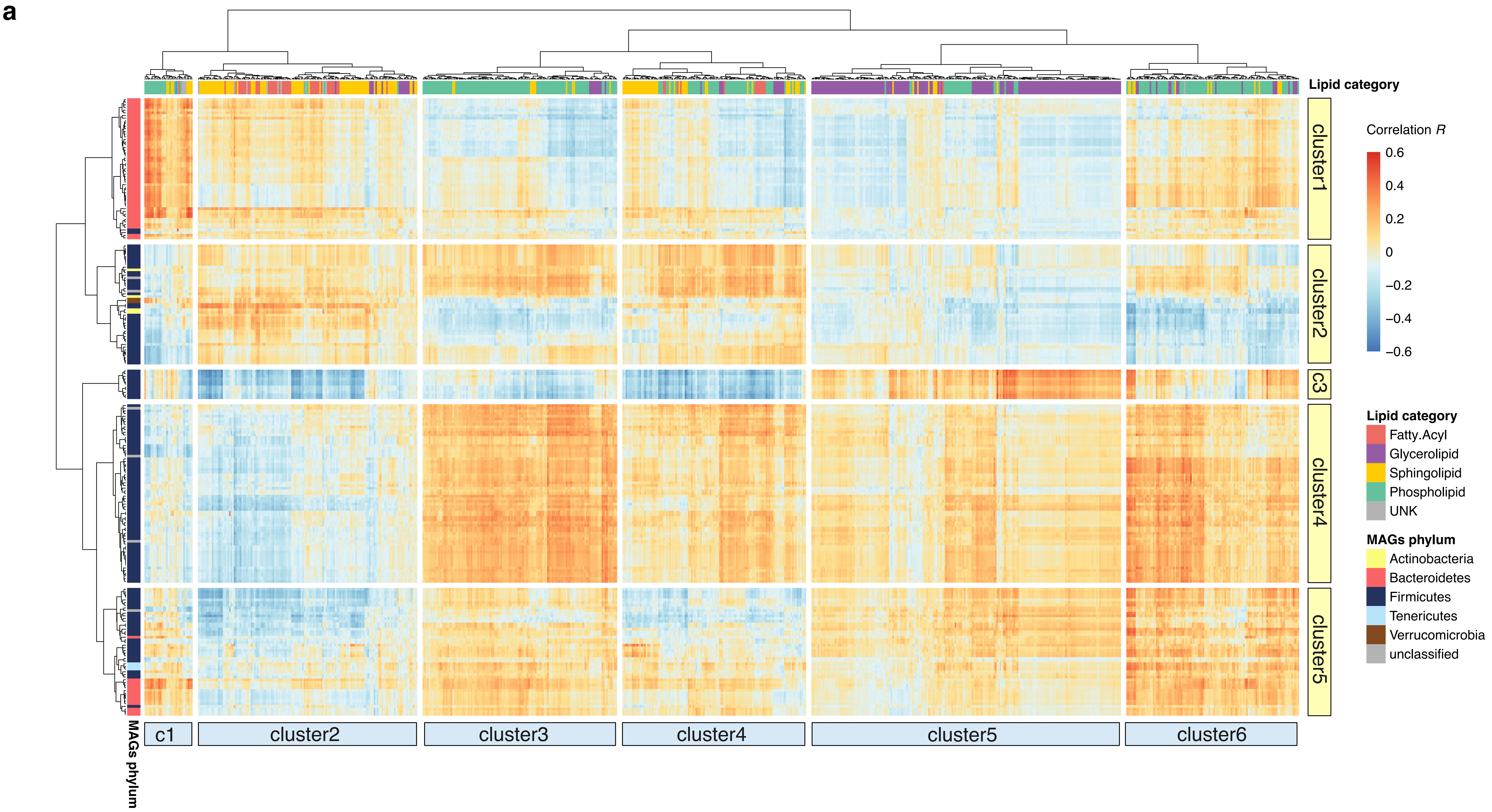


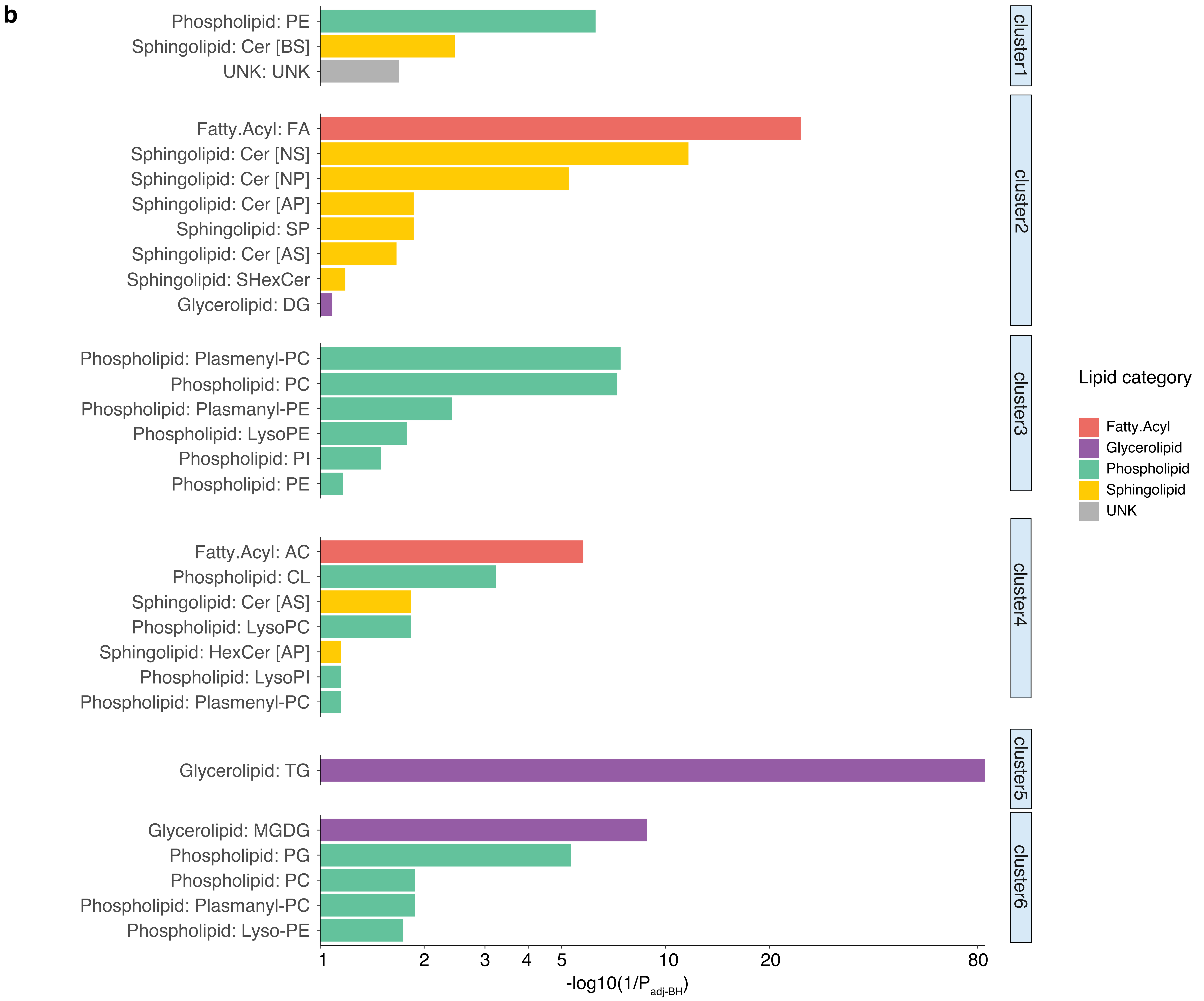

Supplementary Figure 4

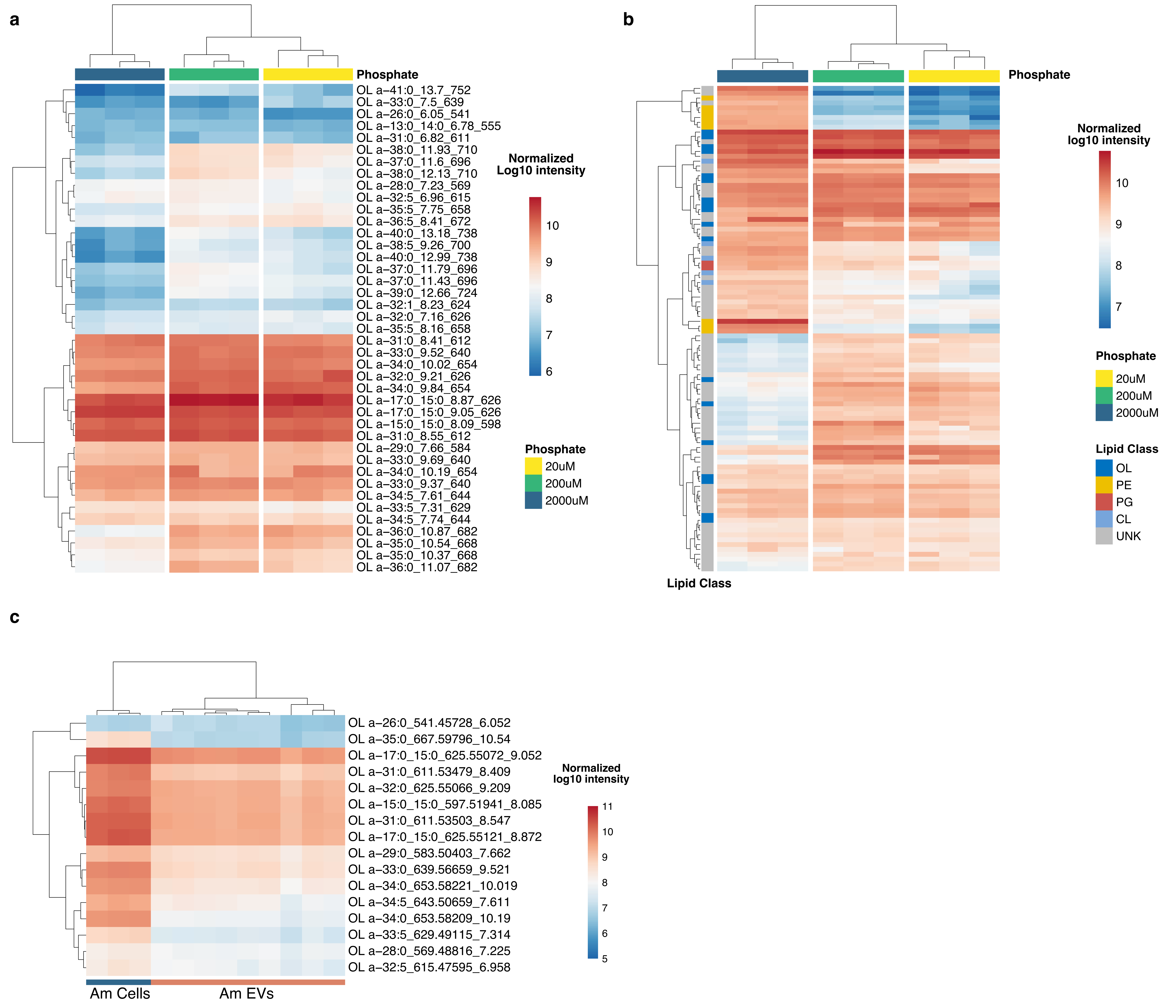

Supplementary Figure 5

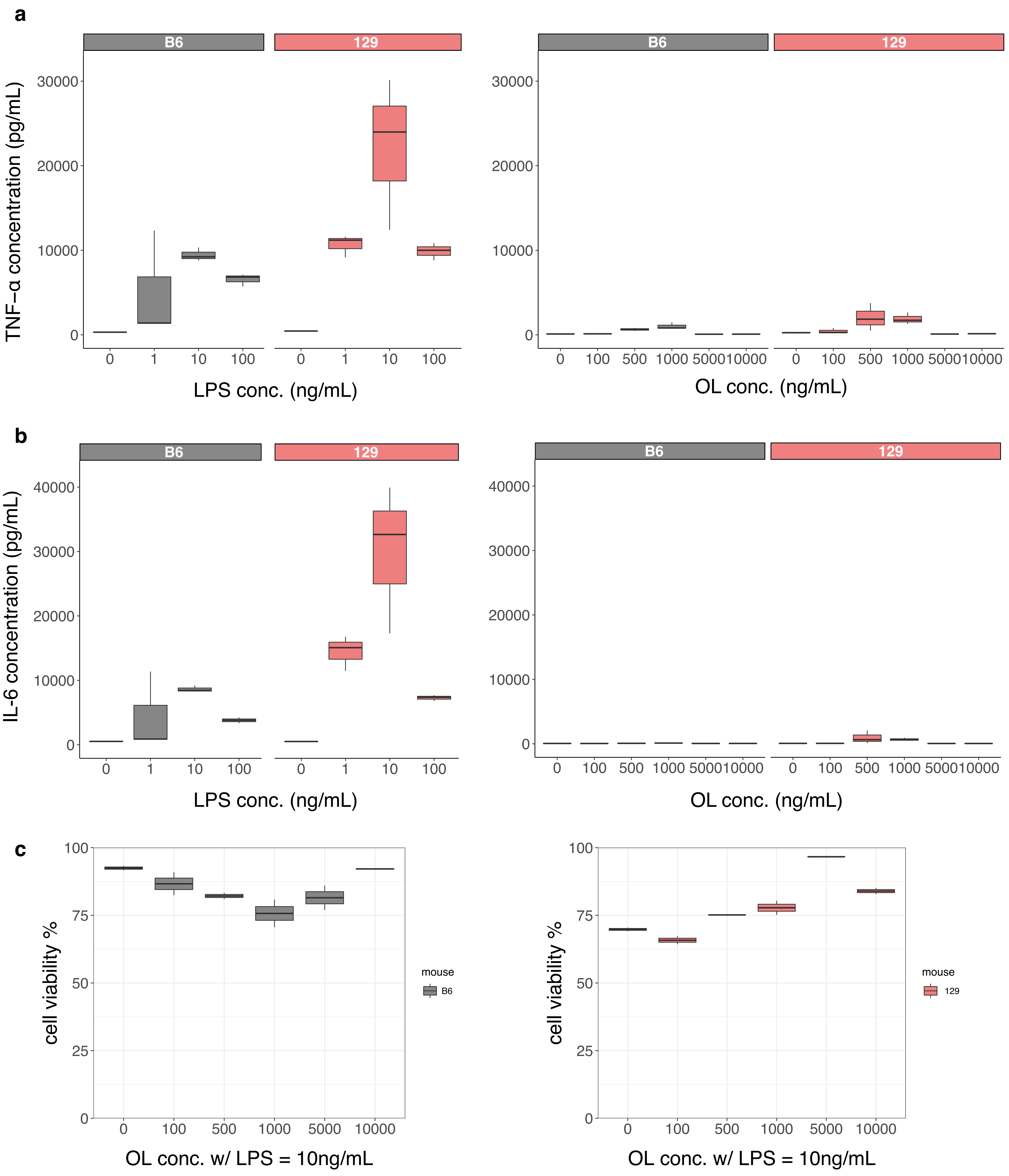

### Supplementary Figure 6

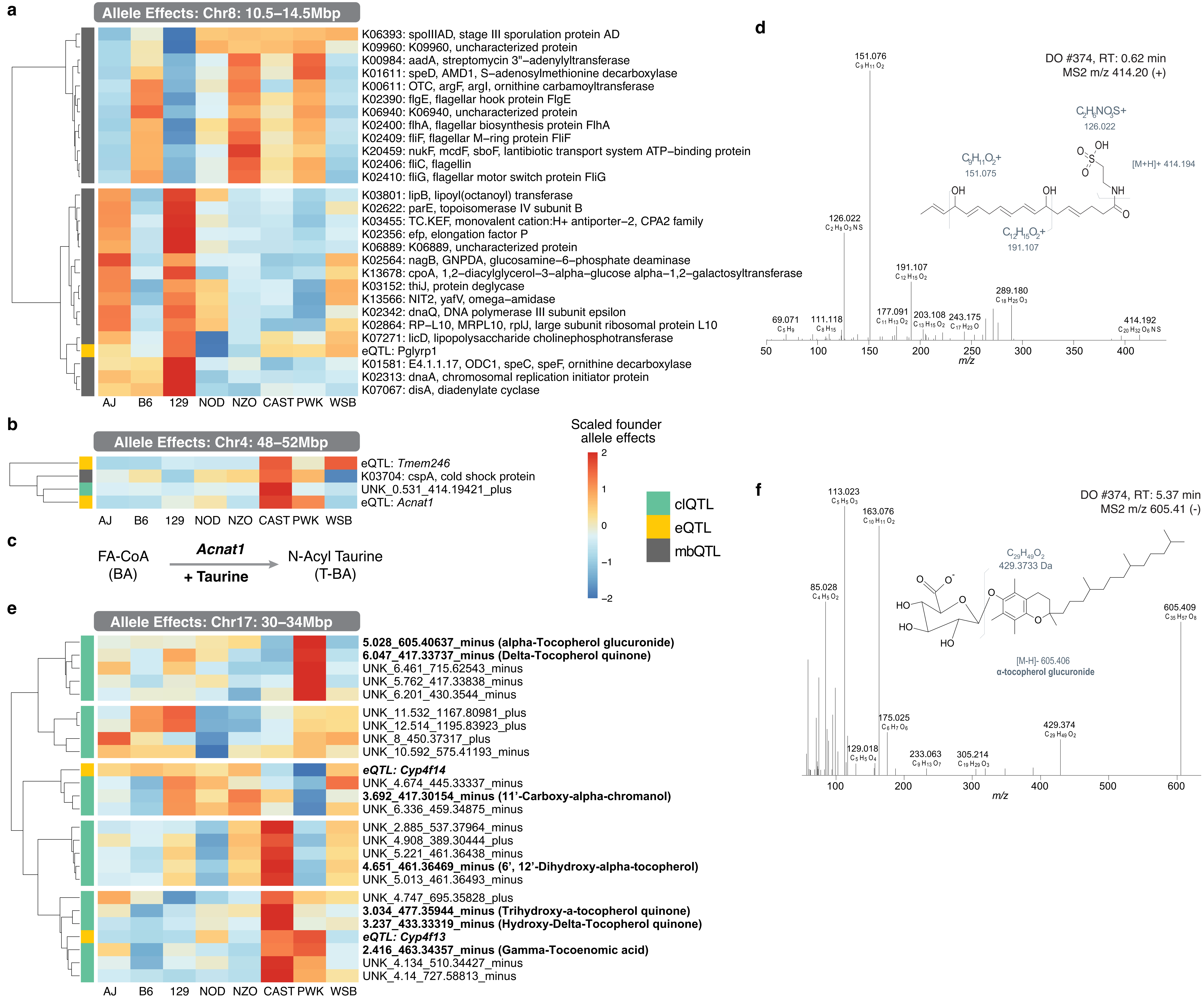

Supplementary Figure 7

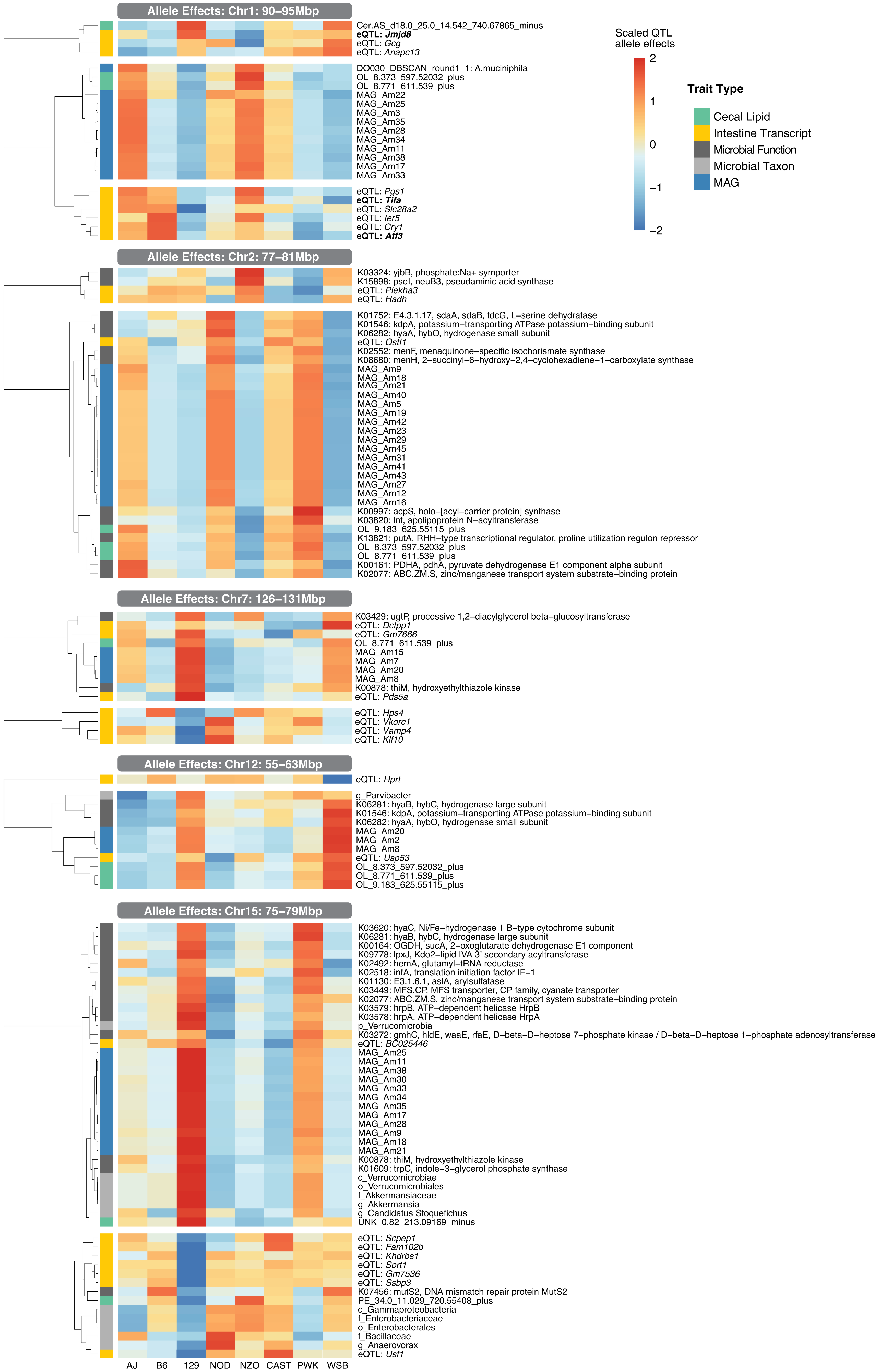
